# Nanopore sequencing reveals operon-specific ribosome remodeling accompanying naphthyridone resistance in *Staphylococcus aureus*

**DOI:** 10.64898/2025.12.11.693786

**Authors:** Dionysius W. Copoulos, Kelly T. Hughes, Fabienne F. Chevance, Cynthia J. Burrows, Aaron M. Fleming, Ryan E. Looper

## Abstract

Antimicrobial resistance (AMR) threatens global health; however, the molecular adaptations underlying resistance to emerging antibiotic classes remain poorly defined. Here, we applied long-read DNA and direct RNA nanopore sequencing and developed methods to deconvolute operon-specific epitranscriptomic changes. Together, this platform uncovered a previously unrecognized, operon-specific pathway of resistance in *Staphylococcus aureus* to the naphthyridone antibiotic A-692345. Genomic nanopore sequencing identified a single 23S rRNA mutation (T1732C) confined to one of the six rRNA operons (operon 2), which uniquely contains nine tRNA genes. RNA direct nanopore sequencing generated a comprehensive and updated rRNA modification map for *S. aureus* and revealed extensive remodeling of rRNA modifications in the resistant strain. Differentially incorporated modifications included pseudouridine, dihydrouridine, and 5-hydroxycytidine at functionally relevant positions within the ribosome. Upon mapping these epitranscriptomic changes, we noted they were operon specific. This likely gives rise to ribosome heterogeneity with potential for selective translation of stress-response genes that favor resistance. Collectively, these findings establish nanopore sequencing as a powerful platform for resolving coupled genomic and epitranscriptomic adaptations, providing molecular insight into how bacteria evolve resistance to antibiotics.

## Introduction

Antimicrobial resistance (AMR) poses an escalating threat to global public health, with resistant pathogens driving increased morbidity, mortality, and healthcare costs worldwide.^1,2^ Bacteria acquire AMR through spontaneous genomic mutations or horizontal gene transfer of mobile resistance elements, enabling the emergence of diverse antibiotic resistance mechanisms.^3^ These mechanisms include enzymatic inactivation of antibiotics, activation of efflux pumps to expel the drug, altered membrane permeability to limit drug uptake, metabolic bypass pathways to circumvent the inhibited process, or enzymatic modification of the antibiotic target to prevent drug binding. High-throughput, short-read DNA sequencing has been instrumental in delineating resistance pathways at the genomic level; however, this approach cannot fully resolve complex genomic regions, large mobile elements, or repetitive sequences, leading to an incomplete understanding of resistance mechanisms.^4^ Long-read nanopore DNA sequencing now provides the accuracy needed to overcome these limitations, while direct RNA nanopore sequencing uniquely enables detection of epitranscriptomic modifications.^4,5^ Here, we employ both DNA and RNA nanopore sequencing to reveal a previously uncharacterized pathway of resistance in *Staphylococcus aureus* (*S. aureus; Sa*) to naphthyridone antibiotics.

The naphthyridone antibiotic class, which was originally described in 2003, has two members, **A-72310** and **A-692345**, that are a promising but underexplored class of antimicrobial agents (Figure 1A).^6^ These compounds display broad-spectrum activity against both Gram-positive and Gram-negative pathogens and selectively inhibit bacterial, but not eukaryotic, translation. Notably, they do not exhibit cross-resistance with established ribosome-targeting antibiotics such as macrolides, tetracyclines or aminoglycosides, indicating a distinct mechanism of action. Initial biochemical analyses implicated the decoding center of the 30S subunit as the target, whereas more recent structural studies challenge this interpretation.^7,8,9^ It was also shown that these naphthyridones can inhibit both the bacterial ribosome and DNA gyrase, suggesting a dual mechanism that may contribute to their broad-spectrum efficacy.^7^ Despite their structural similarities to the quinolone antibiotics, they retain potency against quinolone-resistant strains suggesting that they may target both bacterial transcription and translation by unique mechanisms. Despite these intriguing features, the precise inhibitory mechanism of bacterial transcription and translation by the naphthyridones remains unknown and they have not been further developed, likely reflecting uncertainties surrounding their multi-target behavior and complex structure–activity relationships. These compounds represent an intriguing tool to uncover both new mechanisms of antimicrobial resistance development and potential new mechanisms of action to combat AMR with new antibiotics.

**Figure 1.**
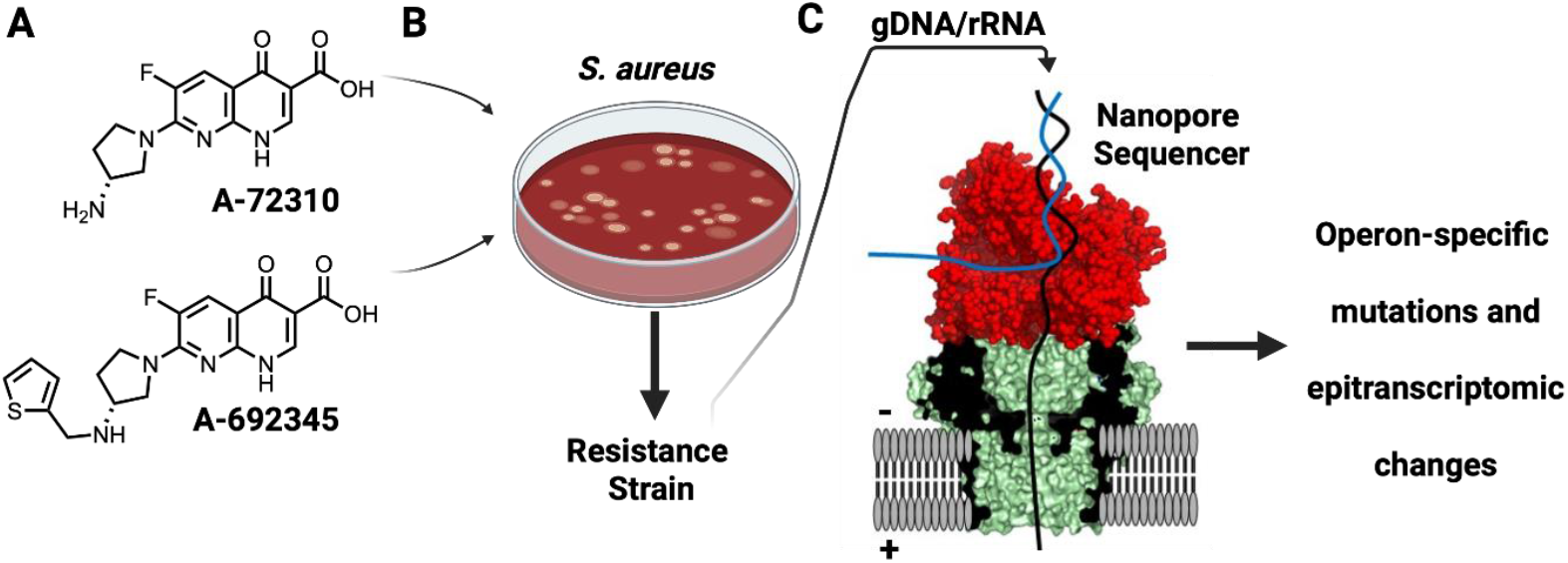
*S. aureus* exposed to naphthyridones yields a resistant strain in which nanopore sequencing reveals operon-specific epitranscriptomic changes. (A) Chemical structures of the naphthyridone antibiotics **A-72310 and A-692345.**(B) *S. aureus* cultures were exposed to the naphthyridones to generate resistant strains. (C) Nanopore sequencing of genomic DNA and rRNA identifies operon-specific mutations and epitranscriptomic changes in the resistance strain.

Extensive investigation into resistance mechanisms in *S. aureus* to antibiotics is known;^10,11^ however, it is not known how this pathogen might adapt to antibiotic stress exerted by these ribosome targeting naphthyridone antibiotics (Figure 1B). Previous work on AMR has focused on acquired resistance through chromosomal mutations and plasmid-borne gene transfer, leaving unexplored the possibility of coordinated genomic mutations and epitranscriptomic responses that facilitate adaptation. The ability of nanopore sequencing to resolve complex, repetitive genomic regions and detect epitranscriptomic modifications provides an opportunity to capture these multilayered antibiotic adaptations in unprecedented detail.^12,13,14^ To address this gap, we combined long-read DNA and direct RNA nanopore sequencing to define the genomic and transcriptomic basis of *S. aureus* resistance to naphthyridone antibiotics (Figure 1C). This approach uncovered operon-specific epitranscriptomic modifications associated with drug adaptation, identifying a previously unrecognized pathway that links genomic plasticity and operon-level RNA regulation to antimicrobial resistance.

## Results and Discussion

### Antibacterial Activity and Resistance Selection

The (*R*)-enantiomers of **A-72310** and **A-692345** were synthesized with minor modifications to previously described methods (see Supporting Information).^15,16^ Their antibacterial activity was evaluated against *S. aureus* (ATCC® 12600™; wild type; Table 1). Consistent with prior reports, both compounds displayed modest antimicrobial activity, with the thiophene analog **A-692345** exhibiting the greatest potency (MIC = 8 µg/mL) vs **A-72310** (MIC = 128 µg/mL).^6,7,15^ To assess the potential for resistance development, *S. aureus* cultures were exposed to increasing concentrations of the compounds (100x → 400x MIC) on solid media. Resistance emerged only in cultures treated with **A-692345**, yielding a resistant strain designated *Sa*_A-692345^R^. A four-fold increase in MIC to **A-692345** was demonstrated for this strain (8 → 64 µg/mL). No resistant colonies were observed upon exposure to **A-72310**, however *Sa*_A-692345^R^ appeared to be cross-resistant to this compound (MIC >128 µg/mL suggesting a shared mechanism of action. Genomic DNA (gDNA) from *Sa*_A-692345^R^ was subsequently sequenced to identify resistance-associated mutations and elucidate the adaptive response to **A-692345** antibiotic stress.

### Sequencing of gDNA reveals a resistance mutation in an rDNA operon in the *Sa*_A692345^**R**^ **strain**

Genomic DNA from *S. aureus* WT and the A-692345^R^ strains was subjected to high-throughput, short-read sequencing, and sequence variants (i.e., mutations) unique to the resistant strain were identified using the GATK pipeline.^17^ A notable mutation was detected in the 23S rDNA gene at position 1732, where a T→C substitution occurred with an allele frequency of ∼17% based on this sequencing approach (Figures S1 and S2). This observation suggests that **A-692345** targets the large ribosomal subunit. Furthermore, the absence of mutations in DNA gyrase or known drug transporters argues against these pathways directly contributing to resistance.

The ∼17% mutation frequency of the 23S T1732C variant is intriguing, as *S. aureus* contains six rRNA operons (1/6 ∼ 17%; Figures 2A and 2B), implying that the mutation resides in only one operon, and that it must be a dominant allele. Inspection of the short-read data supported this interpretation but could not definitively assign the mutation to a specific operon (Figure 2C), due to the inability of short-read sequencing to uniquely map reads within the long repetitive rDNA regions. To overcome this limitation, we performed long-read nanopore DNA sequencing. Reads exceeding 5 kb were mapped to the *S. aureus* genome, revealing that a single operon, operon 2, harbored the 23S T1732C mutation at 100% frequency (Figures 2B and D). Together, these findings demonstrate that resistance to **A-692345** arises from a single rDNA operon-specific mutation in the 23S rDNA, raising questions about how ribosomal heterogeneity at a single operon contributes to naphthyridone resistance.

**Figure 2.**
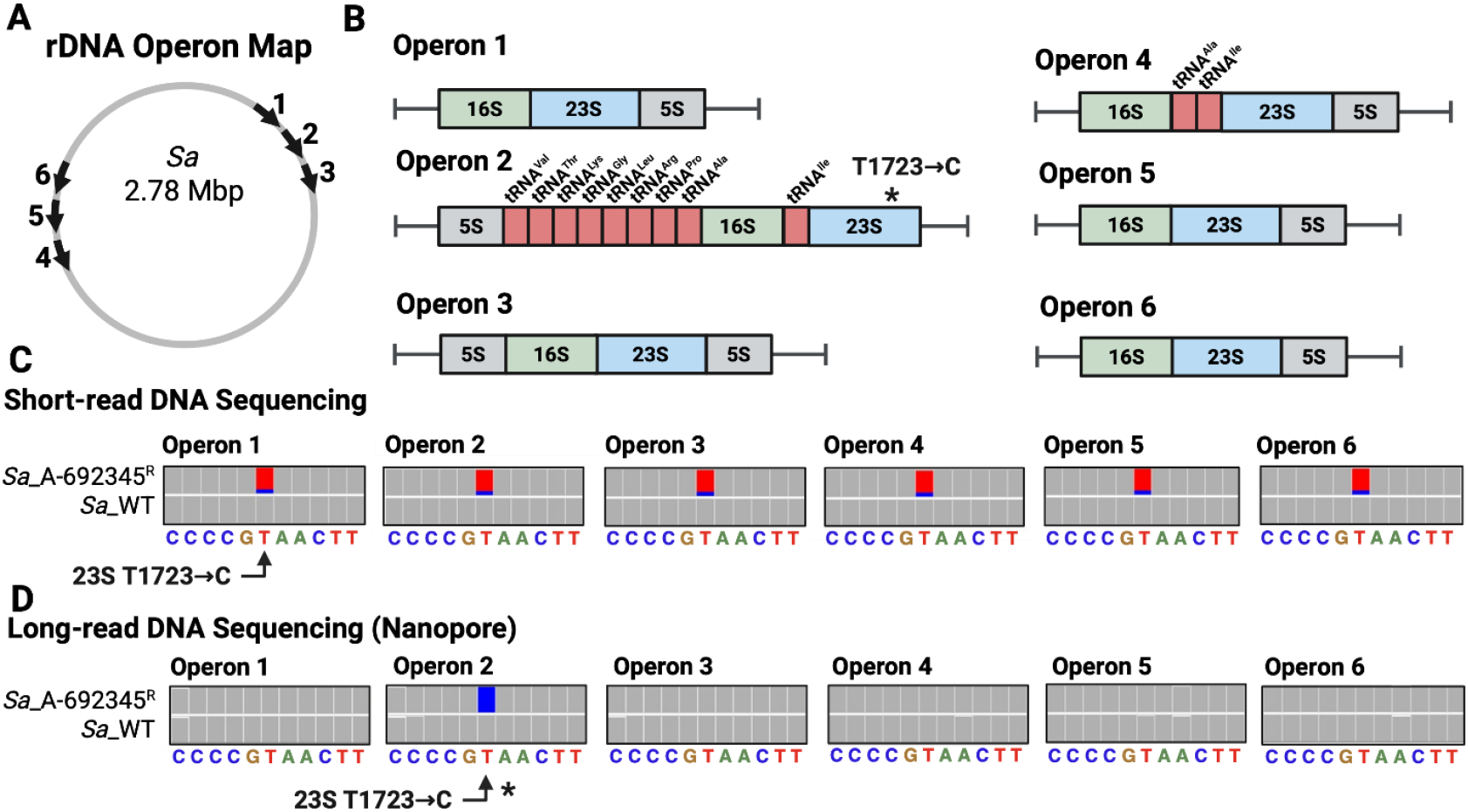
The *S. aureus* genome contains six rRNA operons, one of which harbors the mutation identified in the A-692345^R^ strain. (A) Map of the six *S. aureus* rRNA operons. (B) Gene organization within each operon. (C) Alignment of short-read DNA sequencing data to the *S. aureus* reference genome showing the 23S rRNA T1732C mutation at ∼17% frequency across all six operons. (D) Alignment of long-read DNA sequencing data to the *S. aureus* reference genome reveals that the 23S rRNA T1732C mutation is present exclusively and with full penetrance in operon 2.

These genomic data from the *Sa*_A-692345^R^ strain provide intriguing insights into the early stages of resistance development. The retention of all six rDNA operons in the resistant strain is notable, as operon loss has been reported in other *S. aureus* resistance contexts, such as MRSA.^11,18^ This operon preservation supports a model in which resistance arises through targeted, operon-specific adaptation rather than global genomic reduction of rDNA operons. The identification of the 23S T1732C mutation exclusively in operon 2 is particularly interesting, as this operon is the longest of the six and contains nine tRNA genes (Figure 2B). Such extended operon architecture is not arbitrary, as it enables coordinated transcription of rRNA and tRNA genes to balance ribosome biogenesis with tRNA supply.^19,20^ The selection of a mutated rDNA operon sequence under antibiotic pressure introduces heterogeneity into the ribosome pool, a phenomenon previously observed in *E. coli* to modulate stress-response gene expression and cellular phenotype.^21^ It is noteworthy that all operons were retained and only one carried the mutation following short-term A-692345 exposure; with prolonged selection, rDNA operon reduction or gene conversion might occur, potentially propagating the mutant allele to additional operons. Mutant propagation to other rDNA operons has been noted in *S. aureus* under linezolid stress as an approach to confer greater antibiotic resistance,^11,18^ which may occur with extended **A-692345** exposure. Nonetheless, these long-read nanopore sequencing data reveal a previously undocumented mode of operon-specific divergence as a potential resistance mechanism, wherein the consequences would be reflected in the rRNA transcript pool.

In the genome, the 23S T1732C mutation is formally a d(T:A) → d(C:G) base pair transition mutation, which yields clues as to the origin of chemistry driving the selected mutant. The d(T:A) → d(C:G) transition originates from dA deamination to 2′-deoxyinosine (dI), an isocoder of dG, which upon replication locks in the d(C:G) mutation.^22^ Nitrosative stress drives dA → dI deamination.^23^ Interestingly, *S. aureus* expresses an antibiotic-inducible nitric oxide synthase that elevates the reactive nitrogen species production responsible for dA deamination.^24^ Thus, a plausible mechanism is that **A-692345** exposure induces *S. aureus* nitric oxide synthase to drive this mutation profile, which would occur most readily in actively transcribed regions of the genome, such as rRNA operon 2, housing many essential tRNA genes.

### RNA direct nanopore sequencing provides a comprehensive map of rRNA epitranscriptome in *S. aureus*

Next, we examined how rRNA heterogeneity and modification patterns differ between the WT-strain *Sa*_ATCC-12600 and *Sa*_A-692345^R^ strains. To accomplish this, we first established an rRNA modification map for *S. aureus*, as the two available cryo-EM structures of the *S. aureus* ribosome lack comprehensive modification annotations, most notably omitting pseudouridine (Ψ) sites.^25,26^ RNA direct nanopore sequencing provides a unique capability to directly detect and semi-quantify RNA modifications in native molecules.^12,13,27,28,29^ We performed a comparative analysis of RNA direct nanopore sequencing data from *S. aureus* and *E. coli*, the latter serving as a well-characterized reference with 36 documented epitranscriptomic modification sites characterized by MS,^30^ cryo-EM,^31^ and RNA direct nanopore sequencing.^12,13^ This analysis focused on identifying base-calling errors and signal deviations at known modification sites, as well as using modification-aware base-calling models for Ψ and m^6^A (Figures 3A–D). The resulting modification map was then compared with reported cryo-EM data (Figure 3E).^25,26^ Because *S. aureus* and *E. coli* rRNA sequences are not identical, reference alignments were used to identify corresponding positions for inspection in *S. aureus*; consequently, modification site numbering differs between the two organisms (see the x-axis in Figure 3E, where *E. coli* numbering is shown in parentheses, and Figure S3). This approach establishes, to our knowledge, the most comprehensive rRNA modification landscape for *S. aureus* strain 12600™, enabling direct comparison of modification frequency and distribution between the *S. aureus* wild-type and *Sa*_A-692345^R^ strains.

**Figure 3.**
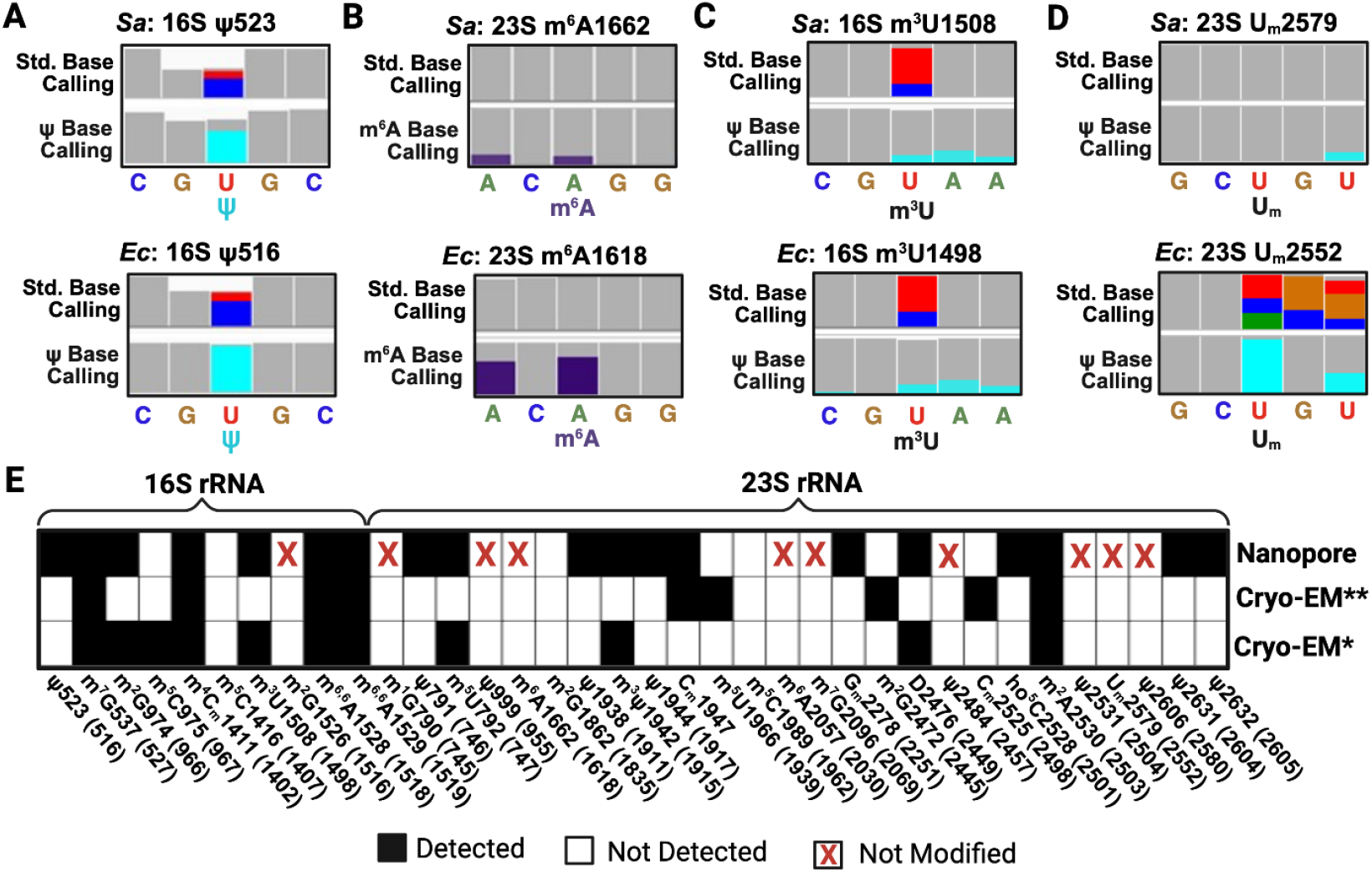
RNA direct nanopore sequencing of native rRNA reveals a comprehensive rRNA epitranscriptomic map in *S. aureus* through comparison with *E. coli* rRNA nanopore data. (A–D) Example Integrative Genomics Viewer (IGV)^32^ analyses showing base miscalls using the standard base call model and modification-aware base calling using modification-aware models at known *E. coli* rRNA modification sites compared with corresponding regions in *S. aureus*. In IGV plots, gray bars indicate correctly aligned sequencing reads to the reference sequence, while base miscalls are color-coded as follows: A = green, C = blue, G = gold, and U = red. Modification-aware base calls are shown as Ψ = turquoise and m6A = purple. (E) Comparison plot of rRNA modification sites identified from RNA direct nanopore sequencing and previously reported cryo-EM data (* = Ref. 26; ** = Ref. 25). Sites identified as modified are marked with a black box; unmodified sites are indicated with a white box. For RNA direct nanopore data, positions showing positive modification signals in *E. coli* but absent in *S. aureus* are marked with a red X to highlight the strong support for the absence of modification at that site. The modification positions are provided for *S. aureus*, and those within parentheses are provided for *E. coli* (Figure S3).

Comparative RNA direct nanopore sequencing revealed a comprehensive rRNA modification landscape in *S. aureus* relative to *E. coli*. Modifications previously reported by cryo-EM in two different *S. aureus* strains (12600™ and RN6390)^25,26^ and confirmed in the nanopore data include 16S m^7^G537, 16S m^2^G974, 16S m^4^C_m_1411, 16S m^3^U1508, 16S m^6,6^A1528, 16S m^6,6^A1529, 23S m^5^U792, 23S m^3^Ψ1942, 23S C_m_1947, 23S D2476, and 23S m^2^A2530 (Figures 3A, 3C, 3E, and S3). Newly identified modifications in *S. aureus*, discovered by comparison to *E. coli* as a benchmark, include 16S Ψ523, 23S Ψ791, 23S Ψ1938, 23S Ψ1944, 23S G_m_2278, 23S ho^5^C2528, 23S Ψ2631, and 23S Ψ2632.

Conversely, several sites that produce strong base-calling signatures in *E. coli* showed no detectable modification signal in *S. aureus* (Figures 3B and 3D), strongly indicating that these positions are unmodified. These sites include 16S G1526 (m^2^G), 23S G790 (m^1^G), 23S U999 (Ψ), 23S A1662 (m^6^A), 23S A2057 (m^6^A), 23S G2096 (m^7^G), 23S U2484 (Ψ), 23S U2531 (Ψ), 23S 2579 (U_m_), and 23S U2606 (Ψ). Consistent with this observation, inspection of the *S. aureus* 12600™ genome (NCBI Ref. Seq: NZ_CP035101.1) did not identify genes encoding the corresponding RNA modification enzymes (“writers”; Figure S3). Given that the *S. aureus* genome (∼2.8 Mbp) is approximately 60% the size of the *E. coli* genome (∼4.6 Mbp), the absence of these genes is not surprising, as many of these modifications, when absent in *E. coli*, do not show strong negative phenotypes.^33,34^

Another modification site in *E. coli* (16S m^2^G1207; Figure S3) was also absent in *S. aureus* because this position corresponds to a cytidine in the latter organism. No additional modification signals were detected elsewhere in the *S. aureus* rRNAs beyond those described above; for example, 23S A2058 in *S. aureus*, when methylated, confers erythromycin resistance.^35^ Three positions observed in the cryo-EM structure of *S. aureus* strain RN6390 were not observed in the present data set on strain 12600™ or the cryo-EM structure of the same strain (Figure 3E; 23S m^5^U1989, 23S m^2^G2474, and 23S C_m_2525); this suggests there may be rRNA modification differences between the strains. These findings validate rRNA modifications previously inferred from cryo-EM^25,26^ and expand the *S. aureus* epitranscriptomic map to include Ψ and ho^5^C. This refined modification map provides a critical foundation for interpreting how naphthyridone antibiotic pressure alters the rRNA modification landscape in the Sa_A692345^R^ resistant strain.

### Operon-specific epitranscriptomic changes in *Sa*_A-692345R

RNA direct nanopore sequencing data can be used to assess the extent of RNA modifications at specific sites. While it is well recognized that nanopore-based modification detection is not as quantitatively accurate as mass spectrometry (MS), it nonetheless provides a powerful comparative approach for identifying changes between samples.^12,13,28^ Because nanopore sequencing errors are largely systematic, they have little impact on relative comparisons between the WT and *Sa*_A-692345^R^ strains. In the WT strain, all six rRNA operons share the same sequence, precluding operon-specific analysis of rRNA modifications. In contrast, in the resistant strain, the 23S rRNA reads can be partitioned based on the U1732C mutation, enabling analysis of operon 2 independently of the other five operons. To ensure that modification signals were specific to the 23S rRNA species containing either the U1732 or C1732 base, sequencing reads were filtered to include only those >2,000 nt in length. This filtering removed short reads that might contain modification signals but not the diagnostic mutation, thus preventing cross-contamination of the operon-specific analyses. This operon-specific separation allows identification of modification changes that may correlate with the unique resistance-associated mutation in operon 2 (23S U1732C).

Comparison of the rRNA modification profiles between the WT and *Sa*_A-692345^R^ strains revealed several changes in both 16S and 23S rRNAs (Figure 4A). For the 16S rRNA, a few sites exhibited fractional modification changes greater than 0.1 (10%). The site showing increased modification in the resistant strain was 16S m^3^U1508 (1.5), whereas decreased modification was observed at 16S m^7^G537 (0.85), 16S m^2^G974 (0.7), 16S m^4^C_m_1411 (0.85), and 16S m^5^C1416 (0.85; Figure 4A).

**Figure 4.**
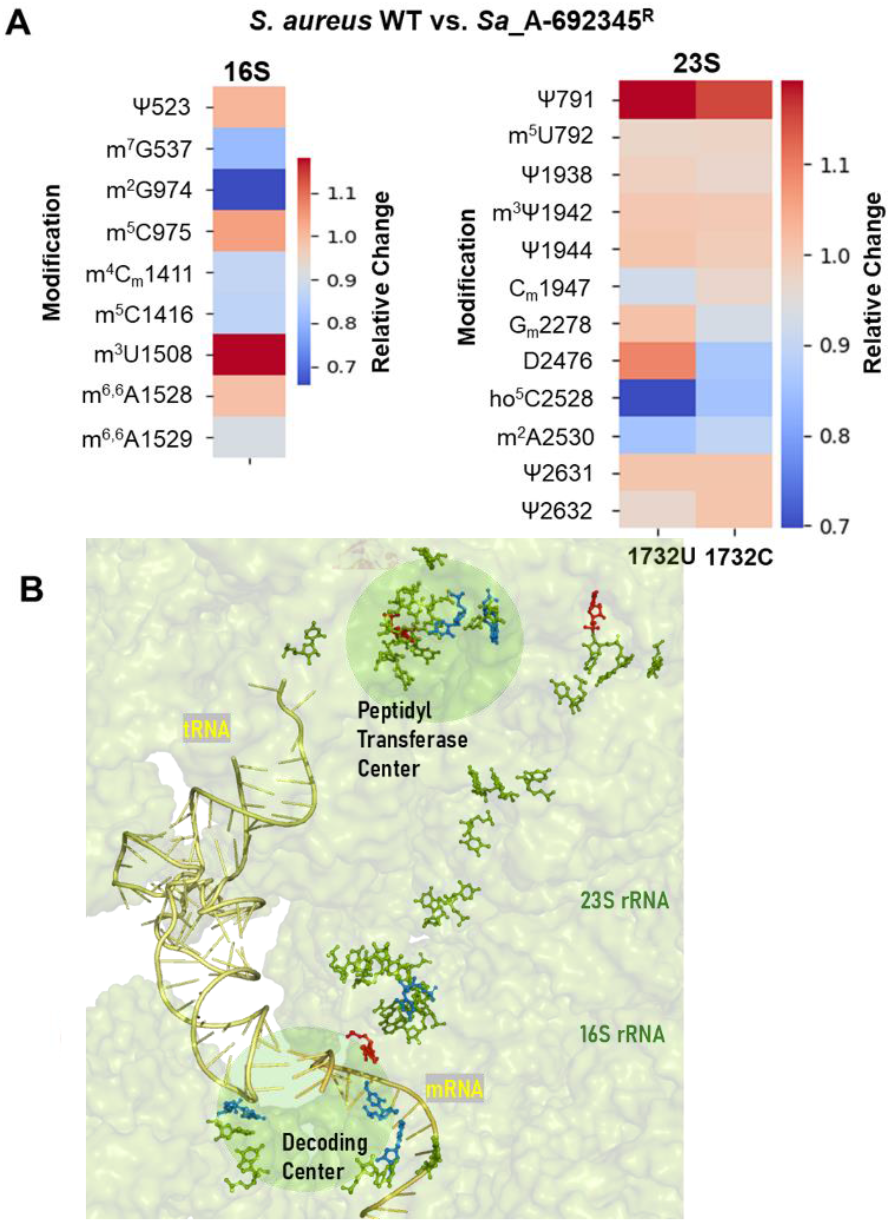
Differentially modified rRNA sites in the *Sa*_A-692345^R^ strain occur in functionally relevant ribosome sites. (A) Heat plots illustrating rRNA epitranscriptomic modification differences between the WT and *Sa*_A-692345^R^ *S. aureus* strains. (B) Ribosome structure displaying the location of modified nucleotides (green), sites of increased modification (red) and decreased modification (blue) relative to functionally relevant centers (pdb 8P2H).^26^

In the 23S rRNA, the expression levels of the U1732 and C1732 sequence variants in the *Sa*_A-692345^R^ strain were similar to those observed in the WT *S. aureus* strain. The key differences in 23S rRNA modification between the WT sequence (U1732) and the mutant (C1732) sequence in the *Sa*_A-692345R strain occurred at positions Ψ791 (U = 1.25; C = 1.1), D2476 (U = 1.05; C = 0.85), and ho^5^C2528 (U = 0.7; C = 0.85; Figure 4A). The D2476 modification in the U1732 sequence increased modestly relative to the WT strain, whereas in the C1732 mutant, the same site showed a pronounced decrease in modification level. For Ψ791 and ho^5^C2528, the U1732 sequence exhibited greater modification changes relative to the WT than did the C1732 sequence. Overall, these results indicate that in the *Sa*_A-692345^R^ strain, the 23S rRNA retaining the native sequence (U1732) underwent more extensive modification remodeling than the mutant (C1732) form. These selective modification changes suggest that naphthyridone exposure induces differential modification control across operons, thereby altering ribosome composition and likely translation dynamics.

Mapping these rRNA modification sites onto a bacterial ribosome structure revealed that nearly all the differentially modified positions localize to functionally important regions, including the peptidyl transferase center and the decoding center (Figure 4B). Changes to these modifications likely influence translation kinetics, fidelity, and/or mRNA selection, providing an additional layer to the resistance phenotype. For instance, 16S m^4^C_m_1411 and m^3^U1508 contact the P-site mRNA codon and fine-tune the geometry of this region to enhance decoding fidelity. Under environmental stress in *E. coli*, decreased methylation at m^4^C_m_1411 has been observed.^12^ In *S. aureus*, the methyltransferases responsible for installing m^4^C_m_ (RsmI and RsmH) contribute to oxidative stress protection;^36^ thus, during **A-692345** exposure, these enzymes may be redirected toward oxidative stress defense, resulting in the altered rRNA modification pattern observed. The decrease in 16S m7G537 modification could also support the Sa_A-692345^R^ resistant phenotype, as loss of this modification confers low-level resistance to streptomycin and neomycin.^37^

In the large subunit, ho^5^C2528 is a late-stage modification that occurs after ribosome assembly, suggesting it does not influence global structure but may modulate translation efficiency or responsiveness under stress conditions, as observed in *E. coli*.^38^ The 23S D2476 modification occurs at a conserved position that requires a pyrimidine for ribosome function; although loss of this D in *E. coli* has no overt phenotype,^39^ the presence of the modification may have a subtle influence on translation dynamics. Together, these observations suggest that ho^5^C and D modifications subtly tune ribosome function to support the resistant phenotype. These support a hypothesis that rRNA modifications can impact ribosome dynamics required for naphthyridone resistance. Detailed structural and biochemical studies are needed to define how each modification contributes to AMR.

The observation of operon-specific epitranscriptomic changes in the rRNA of *S. aureus* conferring resistance to the naphthyridone antibiotic **A-692345** was discovered using nanopore sequencing as the enabling tool. The consequence of this operon-specific remodeling of the rRNA epitranscriptome is the generation of a heterogeneous pool of ribosomes within the cell. Increasing evidence indicates that bacterial ribosome heterogeneity under stress conditions serves as a mechanism for selective translation of stress-response genes.^21^ In the *Sa*_A-692345R strain, such ribosome heterogeneity could enable preferential translation of genes that promote AMR. This is a compelling hypothesis that warrants further study, particularly because the operon harboring the 23S rRNA mutation also contains nine tRNA genes, which could further modulate translational output through altered tRNA stoichiometry.^19^ The presence of mutations in tRNA-rich rDNA operons may represent a generalizable mechanism of resistance; however, this is a speculative hypothesis that needs further study. Nonetheless, these findings establish a foundation for future studies to define a mechanism of action for the inhibition of bacterial translation by **A-692345** and to determine whether ribosome heterogeneity plays a causal role in the resistance phenotype.

## Conclusions

In *S. aureus*, resistance to ribosome-targeting antibiotics typically arises through rRNA sequence mutations or the acquisition of new rRNA modifications that prevent drug binding.^3^ However, traditional approaches such as short-read DNA sequencing, qPCR, and gel-based assays are limited in their ability to resolve operon-specific mutations or detect operon-specific epitranscriptomic changes.^4,11,18^ Building on our prior work demonstrating that RNA direct nanopore sequencing can monitor operon-level stress responses in *E. coli*,^12^ we applied a combined DNA and RNA nanopore sequencing strategy to *S. aureus* exposed to the naphthyridone antibiotic **A-692345**. DNA nanopore sequencing revealed a single mutation in the 23S rDNA of operon 2 (U1732C) in the resistant strain *Sa*_A-692345R, while RNA direct nanopore sequencing uncovered operon-specific remodeling of rRNA modifications associated with this mutation. Together, these data reveal an unrecognized mode of ribosome-targeting antibiotic resistance based on operon-specific epitranscriptomic adaptation. This work establishes nanopore sequencing as a powerful approach for uncovering complex, multilayered mechanisms of AMR and provides a foundation for future structural and biochemical studies to define the **A-692345** binding site and its associated resistance phenotype.

## Associated Content

### Supporting Information

Complete methods, IGV plots, and rRNA modification details.

## Data Availability

All base-called short-read and nanopore data, the Python code for the analysis, and modification data files are available upon request.

## Notes

CJB and AMF have a patent licensed to Electronic BioSciences, and AMF occasionally consults on nucleic acid chemistry for Electronic BioSciences.

## Acknowledgements

This work was supported by the National Institute of General Medical Sciences via grant no. NIH R01AI127724 (REL) and R35 GM145237 (CJB). The support and resources from the Center for High Performance Computing at the University of Utah are gratefully acknowledged. The computational resources used were partially funded by the NIH Shared Instrumentation Grant 1S10OD021644-01A1. The NMR results included in this report were recorded at the David M. Grant Center, a University of Utah Core Facility. Funds for the construction of the Center and the helium recovery system were obtained from the University of Utah and the National Institutes of Health awards 1C06RR017539-01A1 and 3R01GM063540-17W1, respectively. The NMR instruments were purchased with support of the University of Utah and the National Institutes of Health award 1S10OD25241-01. The content is solely the responsibility of the authors and does not necessarily represent the official views of the National Institutes of Health.

