## Supplemental Information for "Nanopore sequencing reveals operon-specific ribosome remodeling accompanying naphthyridone resistance in *Staphylococcus aureus*"

| Item | Page |
|---|---|
| **Materials and Methods** | S2 |
| **Figure S1**. Short-read sequencing mutants found in the *Sa*_A-692345$^R$ strain using the GATK pipeline. | S14 |
| **Figure S2**. Agarose gel electrophoresis analysis of RNA directly sequenced. | S15 |
| **Figure S3**. RNA direct nanopore sequencing data visualized by IGV plots. | S16 |
| **Figure S4**. Comparisons of *S. aureus* and *E. coli* RNA direct nanopore data. | S17 |
| **References** | S29 |

### Materials and Methods

#### Synthesis and Characterization of A-72310 and A-692345

All reagents were obtained from commercial sources and used as received, unless otherwise noted. Reactions run under anhydrous conditions were performed under a positive-pressure of nitrogen ($N_2$) using flame-dried glassware. The tetrahydrofuran (THF), dichloromethane ($CH_2Cl_2$), acetonitrile (MeCN), and *N,N*-dimethylformamide (DMF) were degassed with $N_2$ and passed through activated alumina before being used in any reaction. Methanol (MeOH) was distilled from $CaH_2$ before use. The reactions were monitored by thin-layer chromatography (TLC) and visualized by a dual short/long wave UV lamp and stained with an aqueous solution of potassium permanganate, ninhydrin, bromocresol green, vanillin, and/or dinitrophenylhydrazine to determine when they reached completion. Column chromatography on silica gel was performed using Siliaflash® P60 (40-63 µm). Reversed-phase column chromatography was performed on a Teledyne ISCO Combiflash®, with pre-packed C18 columns. Mass spectra were obtained at the University of Utah on a Waters LCT Premier (ESI/APCI-TOF) for HRMS. $^1$H NMR spectra were recorded at 500 MHz or 400 MHz using Agilent DirectDrive 500, Bruker Neo500, or Inova 400 instruments. Chemical shifts (δ) of proton resonances are reported relative to the deuterated solvent peak (7.27 ppm for $CDCl_3$, 3.31 ppm for MeOH-$d_4$, 2.50 ppm for DMSO-$d_6$, 4.79 ppm for $D_2O$) using the following format: chemical shift (multiplicity [s = singlet, br. s = broad singlet, d = doublet, t = triplet, q = quartet, dd = doublet of doublets, dt = doublet of triplets, td = triplet of doublets, qd = quartet of doublets, m = multiplet], coupling constant(s) (*J* in Hz), integral). $^{13}$C NMR spectra were recorded at 125 MHz using Agilent DirectDrive 500 or Bruker Neo500 instruments. Chemical shifts (δ) of carbon resonances were reported relative to the deuterated solvent peak (77.23 ppm for $CDCl_3$, 49.15 ppm for MeOH-$d_4$, or 39.50 ppm for DMSO-$d_6$).

#### Synthesis Protocols

**General Procedure A**- Displacement of naphthyridone chloride via SNAr

In a solution of acetonitrile (0.3M) was dissolved both the naphthyridone compound **S1** (1.0 equiv.) and amine (1.5 equiv.). This solution was then treated with diisopropylethylamine (3.0 equiv.). The reaction solution was heated to reflux (80 °C) under an inert atmosphere and monitored via TLC. Upon completion, the solution was diluted with water, and the product precipitated out and was collected via filtration. The solid was washed with cold water and dried to furnish a white solid. If a precipitate did not form, the product was extracted from the reaction solution with $CH_2Cl_2$. Followed by washing with 1 M HCl, sat. $NaHCO_3$, and brine. The organic layer was dried with $Na_2SO_4$ and concentrated under reduced pressure to give the title compound.

**General Procedure B**- Deprotection

The protected substrate (1.0 equiv.) was dissolved in trifluoroacetic acid (TFA; 0.1M), and the mixture was warmed to 70 °C. After heating, the mixture became a clear, purple solution. The reaction was monitored every hour via TLC. Upon completion, the mixture was cooled and concentrated under reduced pressure, resulting in a dark purple gum. The solution was dissolved in minimal MeCN and triturated with water. The grey precipitate was filtered and washed with cold water. If titration could not be performed, the compound was purified by column chromatography (5-15% MeOH in $CH_2Cl_2$).

**General Procedure C**- Hydrolysis of the ethyl ester

The ethyl ester (1.0 equiv.) was dissolved in ethanol: water (8:2) (0.1M) and LiOH (10.0 equiv.) was added to the solution, and the reaction mixture was stirred at room temperature (RT). Upon completion, the mixture was diluted in 3 volumes of water and brought to a pH of 3 with 5 M HCl, yielding a white precipitate. If necessary, the solid was then purified with C18 reversed-phase silica cartridges using an automated ISCO purification system. A gradient from 0% to 100% of MeCN in water with 0.1% TFA was used to affect the separation. The resulting product was generally an off white solid.

Preparation of *Ethyl 7-chloro-1-(2,4-dimethoxybenzyl)-6-fluoro-4-oxo-1,4-dihydro-1,8-naphthyridine-3-carboxylate* (**S1**)

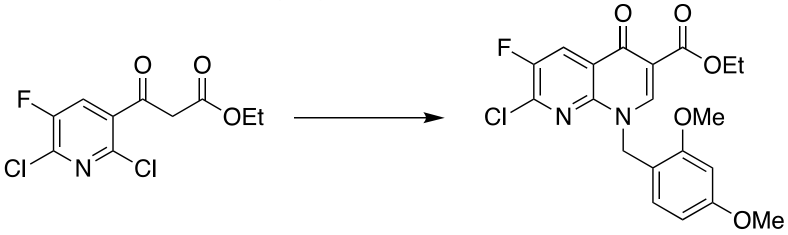


Ethyl 3-[2,6-dichloro-5-fluoro-(3-pyridiyl)]-3-oxopropanoate (107 mmol, 1.0 equiv.) was slurried in acetic anhydride (1.4M) and then treated with triethyl orthoformate (1.2 equiv.). The reaction mixture was refluxed (130 °C) for 14 h. Upon completion, the reaction mixture was concentrated down to afford a brown oil. This brown oil (1.0 equiv.) was then taken up in $CH_2Cl_2$ (0.3 M) and cooled to 0 °C. This solution was treated with 2,4-dimethoxybenzylamine (1.1 equiv.) and stirred at RT for 1 h. Upon completion, the reaction solution was concentrated under reduced pressure. The concentrated product (1.0 equiv.) was dissolved in a solution of MeCN (0.3M) and treated with potassium carbonate (2.0 equiv.). The reaction mixture was heated to reflux (80 °C) under inert conditions for 14 h. Upon completion, the reaction was cooled and extracted with EtOAc. The organic layer was washed with water and 10% aq. citric acid and then dried with $Na_2SO_4$ and concentrated under reduced pressure to give the title compound as a red-orange oil. The crude product was recrystallized with acetone: water (3:1) to furnish a yellow-orange solid. (81%)

$^1$H NMR (400 MHz, $CDCl_3$) δ 8.87 (s, 1H), 8.39 (d, *J* = 7.5 Hz, 1H), 7.46 (d, *J* = 8.3 Hz, 1H), 6.47 – 6.40 (m, 2H), 5.39 (s, 2H), 4.35 (q, *J* = 7.2 Hz, 2H), 3.81 (d, *J* = 1.5 Hz, 3H), 3.76 (d, *J* = 1.5 Hz, 3H), 1.37 (t, *J* = 7.2 Hz, 3H).

$^{13}$C NMR (125 MHz, $CDCl_3$) δ 173.40, 164.75, 161.61, 159.16, 153.24, 151.17, 150.34, 144.55, 141.85, 141.67, 132.96, 124.04, 124.02, 123.27, 123.11, 114.64, 111.25, 104.34, 98.56, 60.92, 55.37, 55.34, 50.75, 14.38.

HRMS calculated for $C_{20}H_{18}ClFN_2O_5$ [M+H] 421.0961; Found 421.0968.

Preparation of *ethyl (R)-7-(3-aminopyrrolidin-1-yl)-1-(2,4-dimethoxybenzyl)-6-fluoro-4-oxo-1,4-dihydro-1,8-naphthyridine-3-carboxylate* (**A-72310a**)

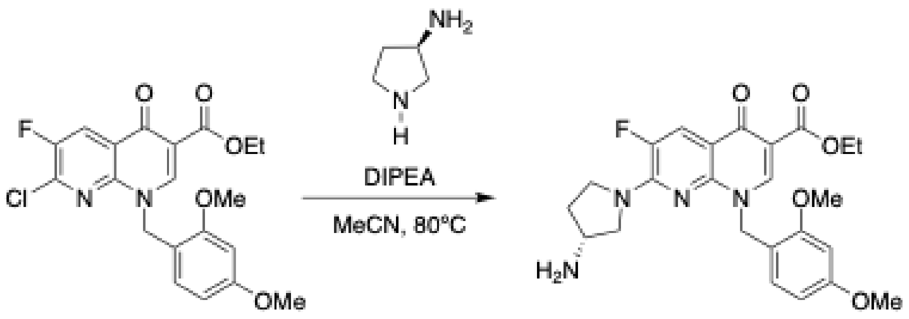

Prepared according to **General Procedure A** from compound **S1** (12.0 mmol). The product was an off-white solid. (99%)
$^1$H NMR (500 MHz, $CDCl_3$) δ 8.59 (s, 1H), 7.88 (d, *J* = 12.9 Hz, 1H), 7.15 (d, *J* = 8.3 Hz, 1H), 6.40 (d, *J* = 2.4 Hz, 1H), 6.37 (dd, *J* = 8.4, 2.4 Hz, 1H), 5.28 (s, 2H), 4.27 (q, *J* = 7.1 Hz, 2H), 3.93 – 3.85 (m, 2H), 3.77 (s, 3H), 3.74 (d, *J* = 2.2 Hz, 1H), 3.72 (s, 3H), 3.71 – 3.67 (m, 1H), 3.50 (ddd, *J* = 11.7, 4.8, 2.6 Hz, 1H), 3.04 (s, 2H), 2.15 (qt, *J* = 7.0, 5.7 Hz, 1H), 1.85 – 1.77 (m, 1H), 1.30 (t, *J* = 7.1 Hz, 3H).
$^{13}$C NMR (125 MHz, $CDCl_3$) δ 174.11, 165.48, 161.21, 158.78, 148.55, 148.46, 148.38, 146.98, 145.39, 144.94, 131.39, 119.37, 119.21, 115.78, 114.42, 114.40, 110.35, 104.29, 98.52, 60.60, 56.51, 56.47, 55.37, 55.29, 50.41, 49.48, 47.21, 47.16, 33.46, 29.65, 14.30.
HRMS calculated for $C_{24}H_{27}FN_4O_5$ HRMS [M+H] 471.2038; Found 471.2038.

Preparation of ethyl *(R)-7-(3-aminopyrrolidin-1-yl)-6-fluoro-4-oxo-1,4-dihydro-1,8-naphthyridine-3-carboxylate* (**A-72310b**)

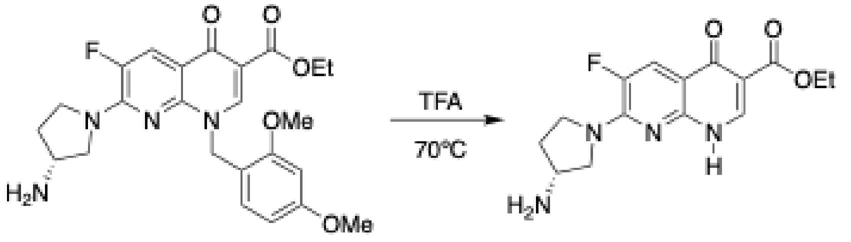

Prepared according to **General Procedure B** from compound **A-72310a** (10.0 mmol). (83%)
$^1$H NMR (500 MHz, DMSO-*d*$_6$) δ 12.47 (s, 1H), 8.45 (s, 2H), 8.30 (s, 1H), 7.69 (d, *J* = 13.0 Hz, 1H), 7.31 (s, 3H), 4.20 (q, *J* = 7.1 Hz, 2H), 3.97 (tt, *J* = 6.0, 3.2 Hz, 1H), 3.85 (ddt, *J* = 32.7, 17.5, 10.6 Hz, 3H), 3.72 (d, *J* = 9.6 Hz, 1H), 2.29 (dp, *J* = 14.7, 7.1 Hz, 1H), 2.16 – 2.08 (m, 1H), 1.27 (t, *J* = 7.1 Hz, 3H).
$^{13}$C NMR (125 MHz, DMSO-*d*$_6$) δ 173.08, 164.45, 158.87, 158.62, 158.37, 158.12, 148.53, 148.43, 146.48, 145.67, 144.46, 144.24, 120.76, 118.38, 118.00, 117.84, 116.00, 113.62, 112.65, 109.48, 59.80, 52.06, 52.01, 49.32, 46.15, 28.67, 14.28.
HRMS calculated for $C_{15}H_{17}FN_4O_3$ HRMS [M+H] 321.1357; Found 321.1358.

Preparation of *(R)-7-(3-aminopyrrolidin-1-yl)-6-fluoro-4-oxo-1,4-dihydro-1,8-naphthyridine-3-carboxylic acid* (**A-72310**)

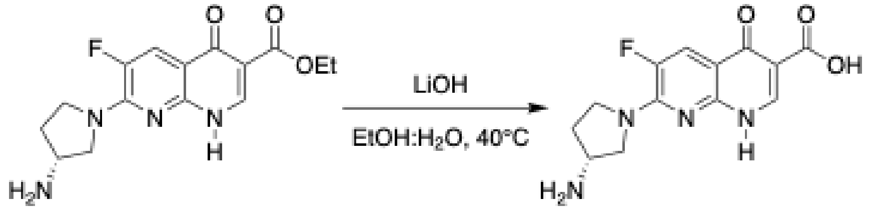


Prepared according to **General Procedure C** from compound **A-72310b** (1.6 mmol). The resulting product was a white solid. (97%)

$^{1}$H NMR (500 MHz, $D_2O$) δ 8.41 (s, 1H), 7.54 (d, *J* = 13.6 Hz, 1H), 3.68 (d, *J* = 17.5 Hz, 2H), 3.58 – 3.46 (m, 2H), 3.28 – 3.21 (m, 1H), 2.12 (dt, *J* = 12.6, 6.4 Hz, 1H), 1.73 (dt, *J* = 12.7, 6.8 Hz, 1H).

$^{13}$C NMR (125 MHz, $D_2O$) δ 181.51, 175.37, 172.96, 166.88, 163.40, 163.12, 162.84, 162.55, 149.02, 148.92, 148.75, 146.76, 144.81, 119.79, 117.48, 116.82, 116.66, 115.16, 113.90, 112.84, 111.66, 55.33, 49.59, 46.82, 38.71, 32.38, 23.27.

HRMS calculated for $C_{13}H_{13}FN_4O_3$ HRMS [M+H] 293.1044; Found 293.1045.

Preparation of *ethyl (R)-6-fluoro-4-oxo-7-(3-((thiophen-2-ylmethyl)amino)pyrrolidin-1-yl)-1,4-dihydro-1,8-naphthyridine-3-carboxylate* (**A-692345a**)

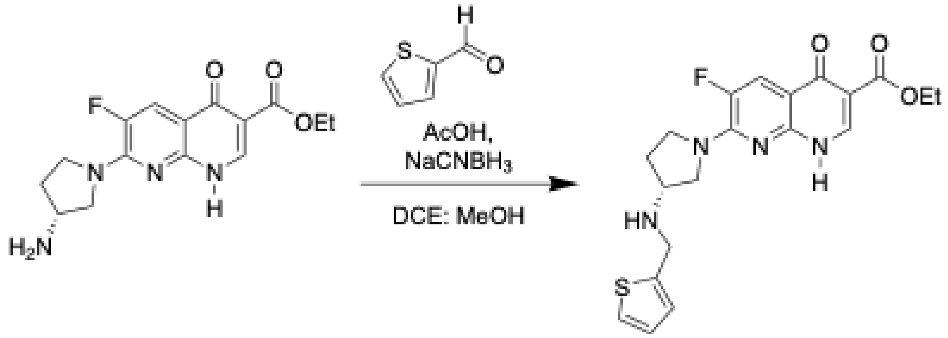


The amine **A-72310b** (0.8 mmol) (1.0 equiv.) and carbonyl compound (1.0 equiv.) were dissolved in DCE (0.1 M) and treated with acetic acid and left to stir for 30 min. Upon consumption of **A-72310b**, the imine was treated with $NaCNBH_3$ (2.0 equiv.). Note: this must be performed in a fume hood, for HCN is a byproduct of this reaction. The reaction was stirred under an $N_2$ atmosphere and monitored via TLC. Upon completion, the reaction was quenched with $NaHCO_3$ and extracted with $CH_2Cl_2$. The organic extract was washed with brine, and the solution was dried over $Na_2SO_4$. The solvent was removed to yield the free base. The compound was then purified by column chromatography (5% MeOH in $CH_2Cl_2$). Note: the aqueous layer should be quenched with bleach for safe waste disposal. (67%)

$^{1}$H NMR (500 MHz, DMSO-$d_6$) δ 8.72 (s, 1H), 7.90 – 7.85 (m, 1H), 7.05 (d, *J* = 8.4 Hz, 1H), 6.58 (d, *J* = 2.3 Hz, 1H), 6.47 (dd, *J* = 8.5, 2.4 Hz, 1H), 5.39 (s, 2H), 3.79 (s, 5H), 3.73 (s, 4H), 3.58 – 3.53 (m, 1H), 3.38 (d, *J* = 12.2 Hz, 1H), 2.05 – 1.97 (m, 1H), 1.76 – 1.66 (m, 2H).

$^{13}$C NMR (125 MHz, DMSO-$d_6$) δ 172.70, 164.54, 148.94, 148.85, 146.63, 146.05, 145.06, 144.61, 143.79, 126.71, 124.65, 124.58, 118.05, 117.89, 112.54, 110.17, 59.67, 54.14, 47.12, 46.00, 14.36.

HRMS calculated for $C_{20}H_{21}FN_4O_3S$ [M+H] 417.1391; Found 417.1392.

Preparation of *(R)-6-fluoro-4-oxo-7-(3-((thiophen-2-ylmethyl)amino)pyrrolidin-1-yl)-1,4-dihydro-1,8-naphthyridine-3-carboxylic acid* (**A-692345**)

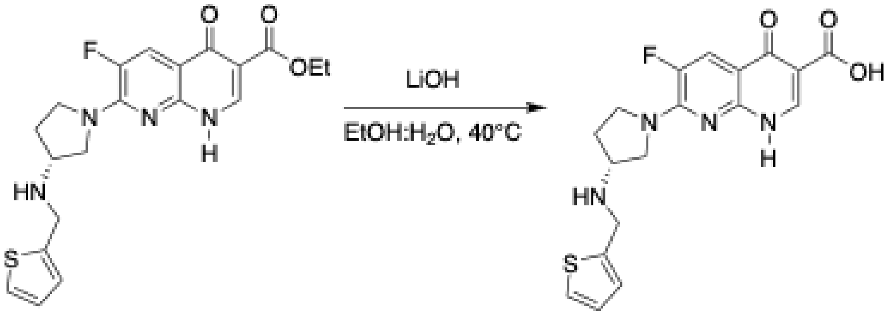


Prepared according to **General Procedure C** from compound **A-692345a** (0.7 mmol). (96%)
$^{1}$H NMR (500 MHz, DMSO-$d_6$) δ 13.30 (s, 1H), 9.38 (d, $J$ = 44.3 Hz, 2H), 8.51 (s, 1H), 8.02 (d, $J$ = 12.8 Hz, 1H), 7.67 (dd, $J$ = 5.1, 1.2 Hz, 1H), 7.33 (d, $J$ = 3.5 Hz, 1H), 7.12 (dd, $J$ = 5.1, 3.5 Hz, 1H), 4.56 (t, $J$ = 11.4 Hz, 2H), 4.09 – 3.93 (m, 5H), 3.83 (s, 2H), 2.42 – 2.29 (m, 2H).
$^{13}$C NMR (125 MHz, DMSO-$d_6$) δ 176.93, 166.05, 149.45, 149.35, 147.19, 146.55, 145.14, 143.70, 132.74, 130.71, 128.50, 127.40, 117.32, 117.16, 110.68, 107.73, 50.68, 46.55, 43.21.
HRMS calculated for $C_{18}H_{17}FN_4O_3S$ [M+H] 389.1078; Found 389.1079.

**Copies of $^{1}$H & $^{13}$C NMR Spectra**

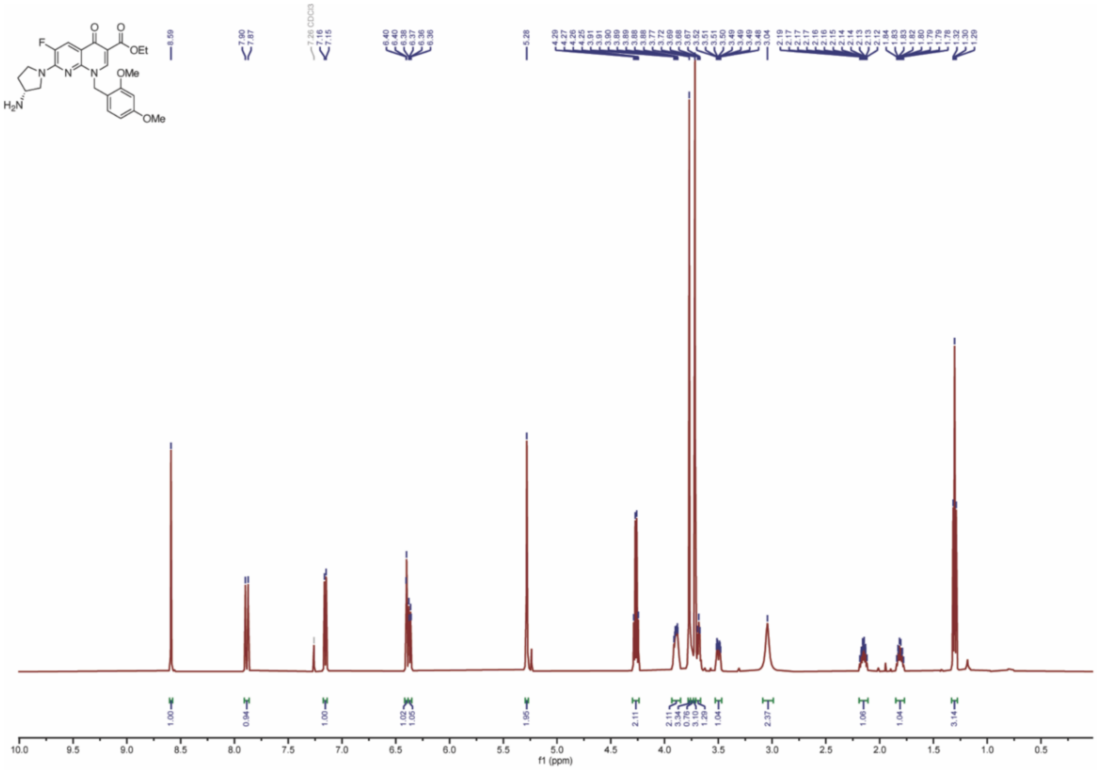
F
O
O
OEt
N
N
N
OMe
OMe
H2N
8.59
7.90
7.87
7.26 CDCl3
7.16
7.15
6.40
6.40
6.38
6.37
6.36
6.36
5.28
4.29
4.27
4.26
4.25
3.91
3.90
3.89
3.89
3.88
3.88
3.77
3.72
3.69
3.68
3.67
3.52
3.51
3.51
3.50
3.49
3.49
3.48
3.04
2.19
2.17
2.17
2.17
2.16
2.16
2.15
2.14
2.13
2.13
2.12
1.84
1.83
1.83
1.82
1.80
1.79
1.78
1.32
1.30
1.29
1.00
0.94
1.00
1.02
1.05
1.95
2.11
2.11
3.34
0.76
3.10
1.29
1.04
2.37
1.06
1.04
3.14
10.0
9.5
9.0
8.5
8.0
7.5
7.0
6.5
6.0
5.5
5.0
4.5
4.0
3.5
3.0
2.5
2.0
1.5
1.0
0.5
f1 (ppm)

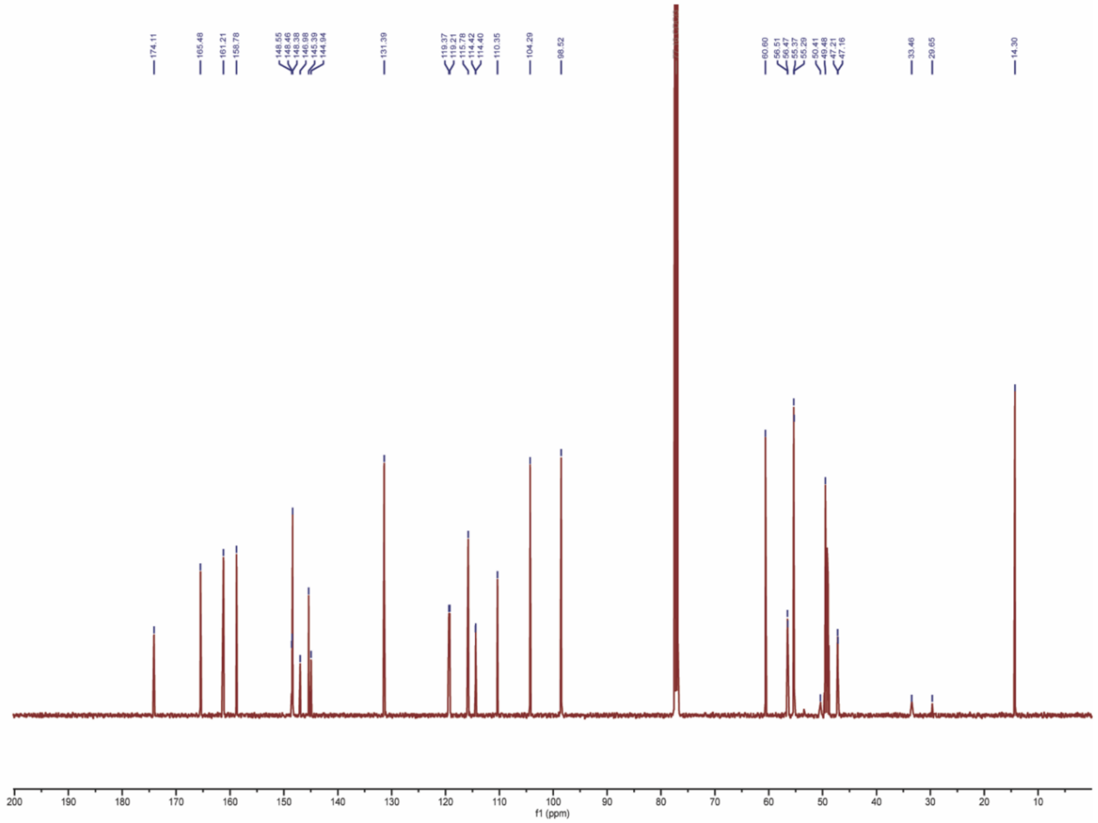
174.11
165.48
161.21
158.78
148.55
148.46
148.38
146.98
145.39
144.94
131.39
119.37
119.21
115.78
114.42
114.40
110.35
104.29
98.52
60.60
56.51
56.47
55.37
55.29
50.41
49.48
47.21
47.16
33.46
29.65
14.30
200
190
180
170
160
150
140
130
120
110
100
90
80
70
60
50
40
30
20
10
f1 (ppm)

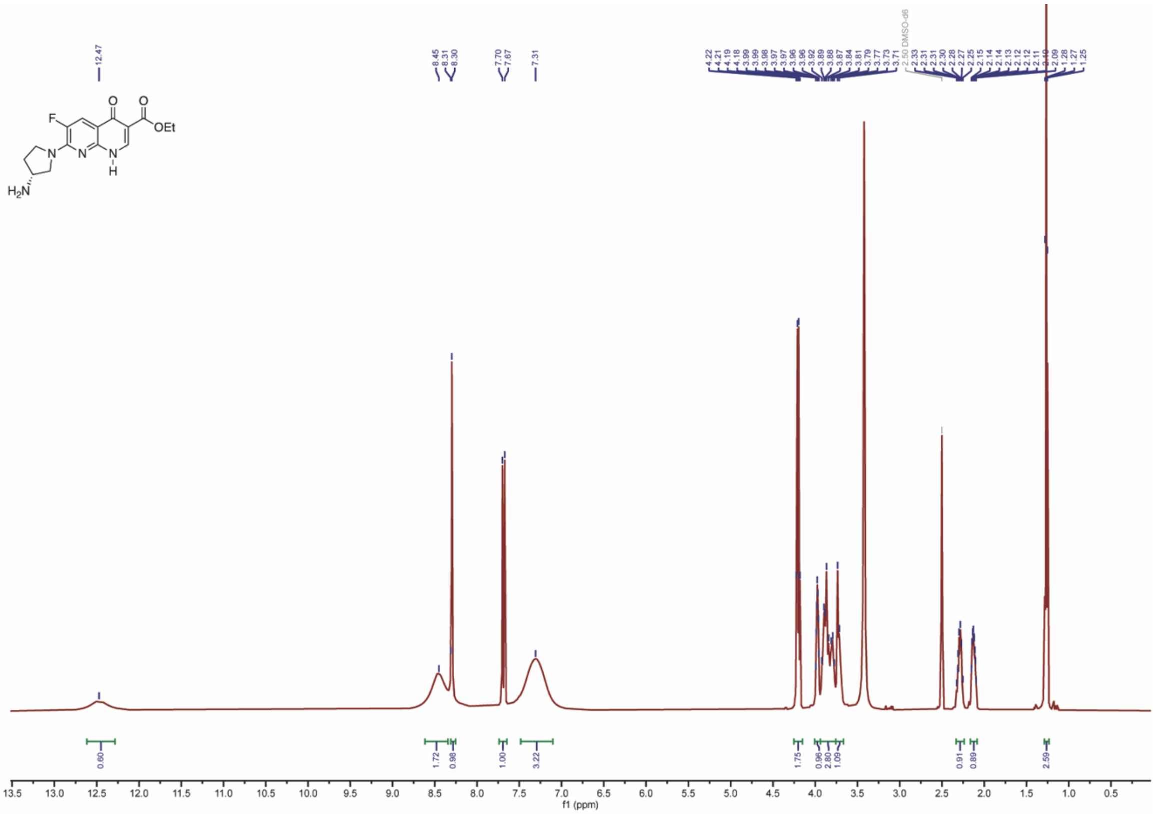

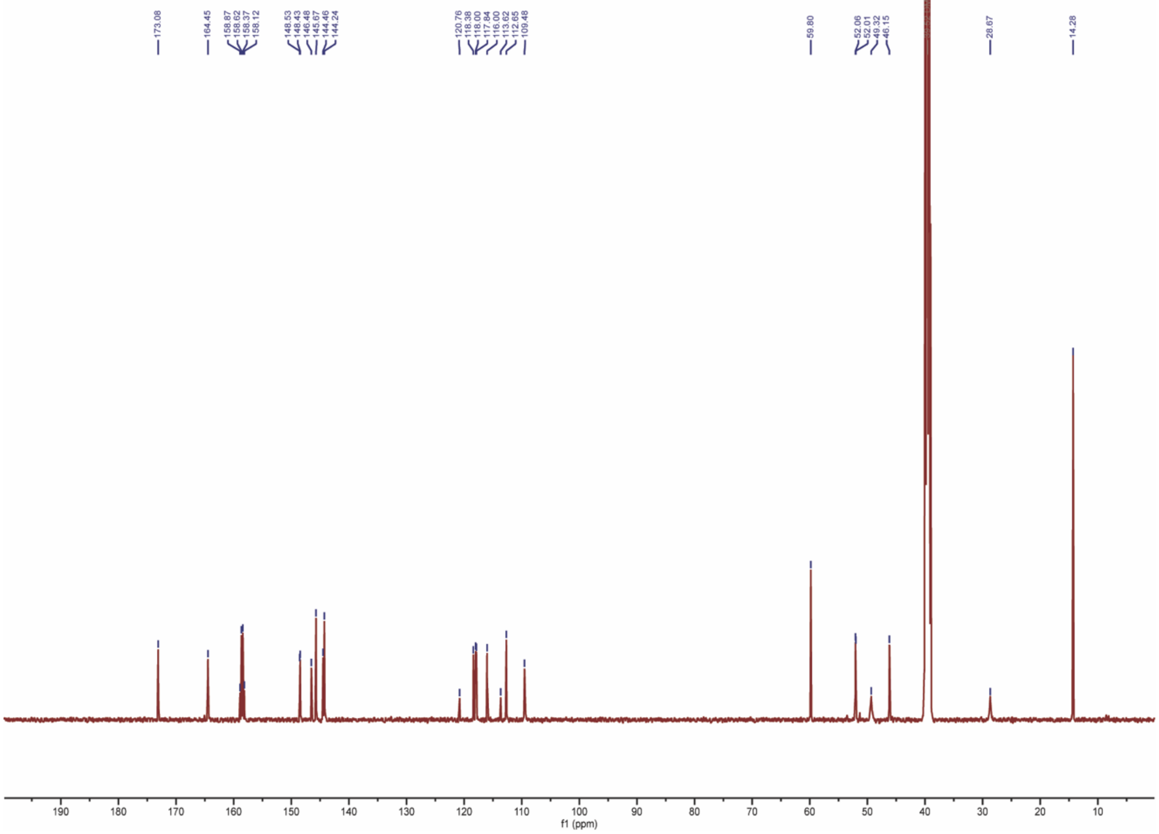

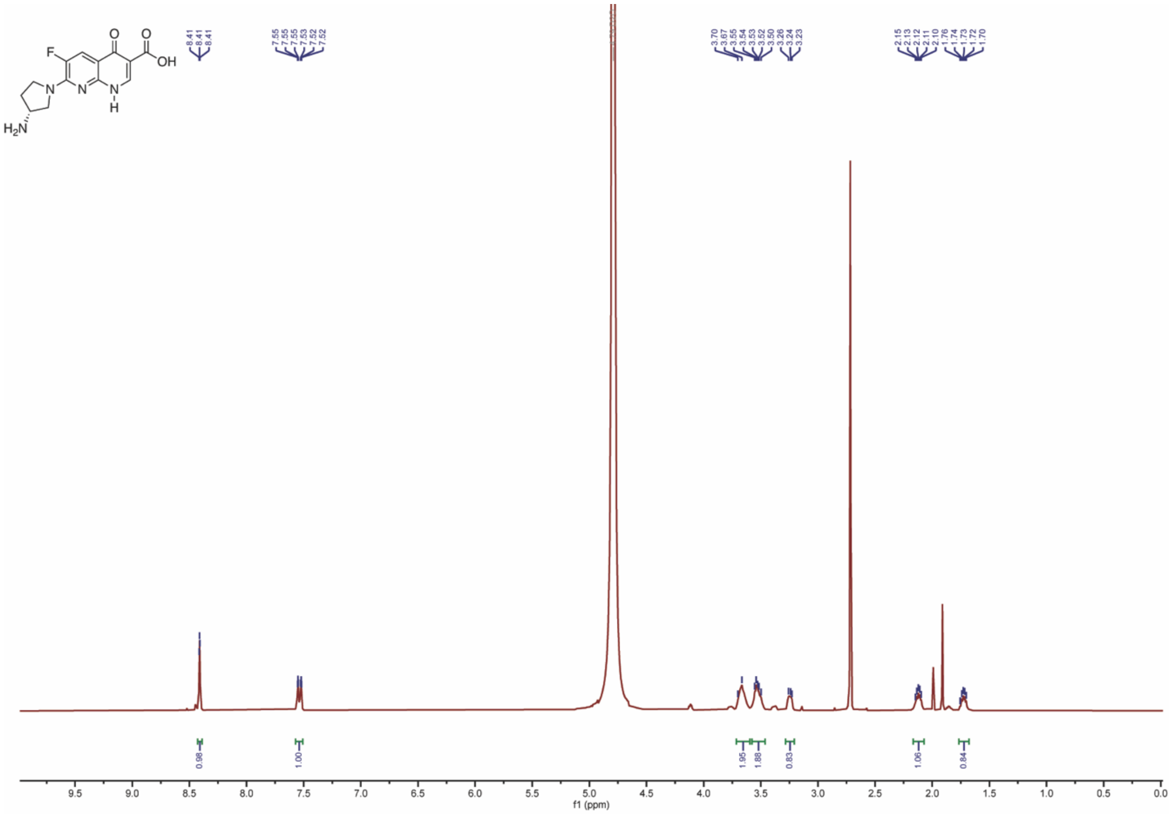
F
O
O
OH
N
N
N
H
$H_2N$
8.41
8.41
8.41
7.55
7.55
7.55
7.53
7.52
7.52
3.70
3.67
3.55
3.54
3.53
3.52
3.50
3.26
3.24
3.23
2.15
2.13
2.12
2.11
2.10
1.76
1.74
1.73
1.72
1.70
0.98
1.00
1.95
1.88
0.83
1.06
0.84
f1 (ppm)

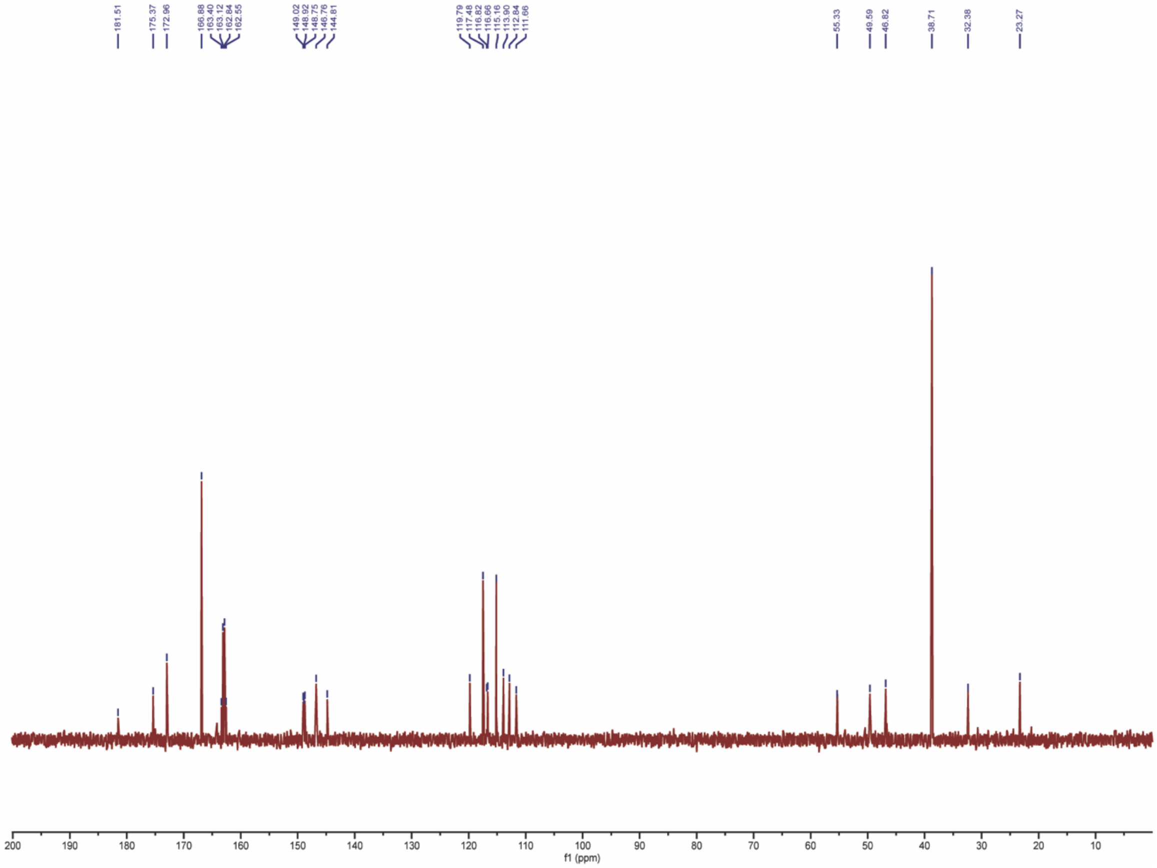
181.51
175.37
172.96
166.88
163.40
163.12
162.84
162.55
149.02
148.92
148.75
146.76
144.81
119.79
117.48
116.82
116.66
115.16
113.90
112.84
111.66
55.33
49.59
46.82
38.71
32.38
23.27
f1 (ppm)

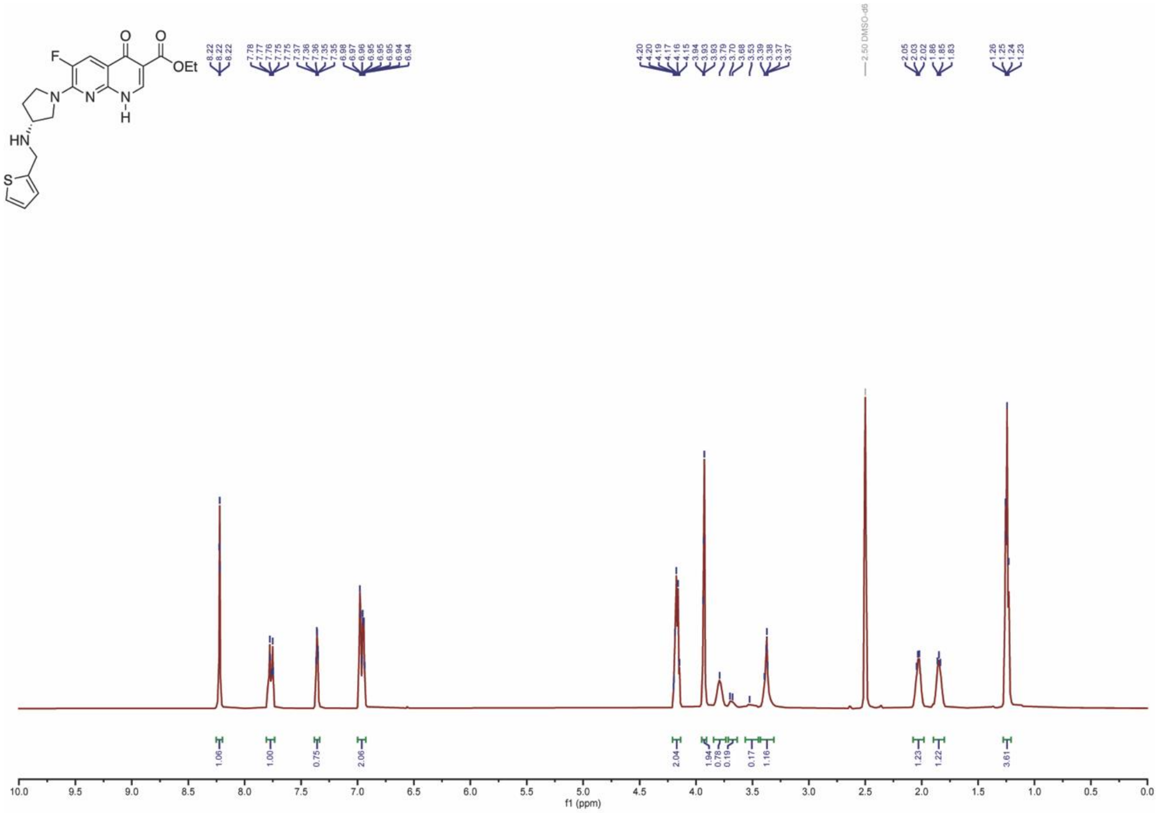
F
O
O
OEt
N
N
N
H
HN
S
8.22
8.22
8.22
7.78
7.77
7.76
7.75
7.75
7.37
7.36
7.36
7.35
7.35
6.98
6.97
6.96
6.95
6.95
6.95
6.94
6.94
4.20
4.20
4.19
4.17
4.16
4.15
3.94
3.93
3.93
3.79
3.70
3.68
3.53
3.39
3.38
3.37
3.37
2.50 DMSO-d6
2.05
2.03
2.02
1.86
1.85
1.83
1.26
1.25
1.24
1.23
1.06
1.00
0.75
2.06
2.04
1.94
0.78
0.19
0.17
1.16
1.23
1.22
3.61
f1 (ppm)

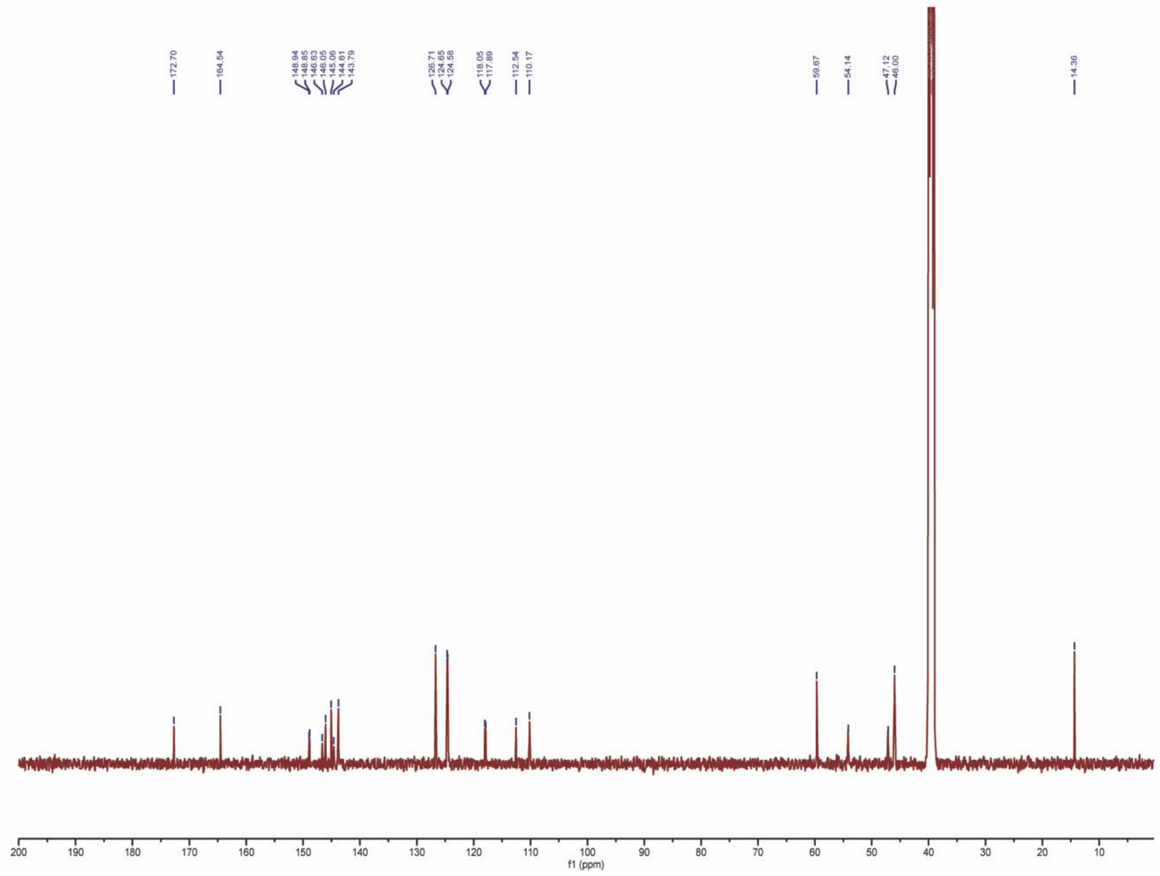
172.70
164.54
148.94
148.85
146.63
146.05
145.06
144.61
143.79
126.71
124.65
124.58
118.05
117.89
112.54
110.17
59.67
54.14
47.12
46.00
14.36
f1 (ppm)

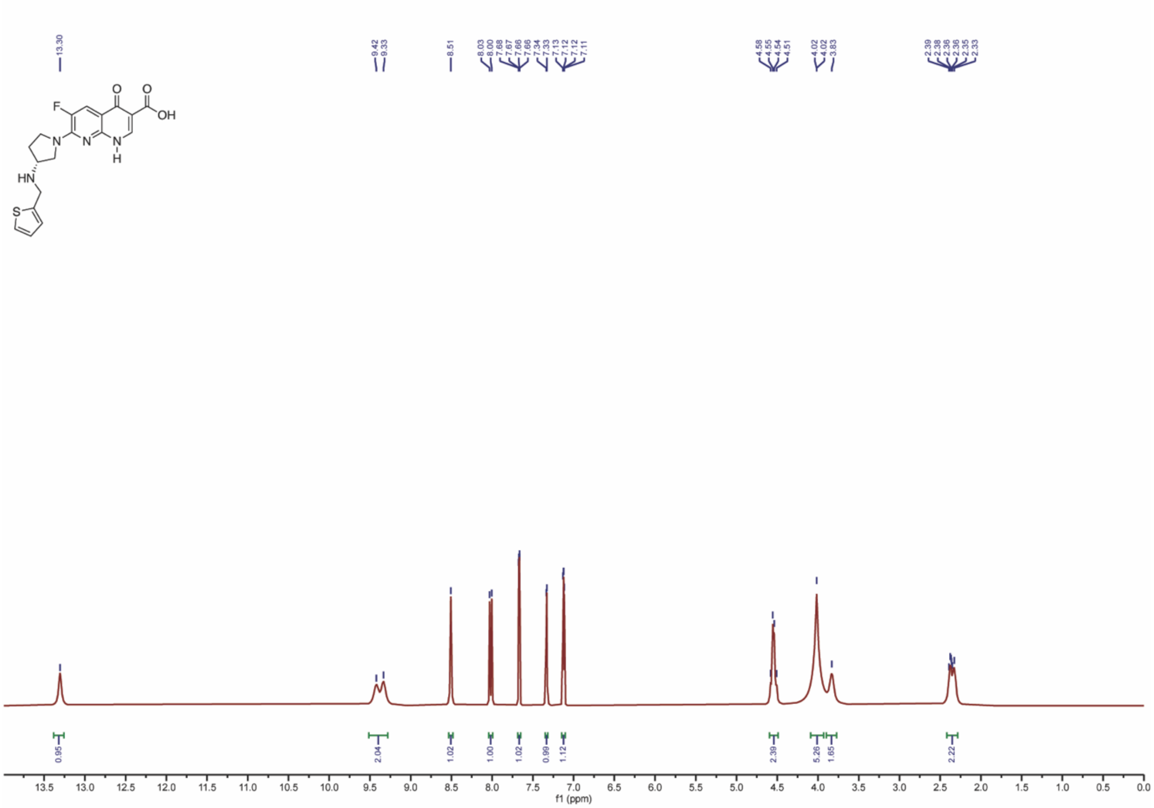
13.30
9.42
9.33
8.51
8.03
8.00
7.68
7.67
7.66
7.66
7.34
7.33
7.13
7.12
7.12
7.11
4.58
4.55
4.54
4.51
4.02
4.02
3.83
2.39
2.38
2.38
2.36
2.35
2.33
F
O
O
OH
N
N
N
H
HN
S
0.95
2.04
1.02
1.00
1.02
0.99
1.12
2.39
5.26
1.65
2.22
f1 (ppm)

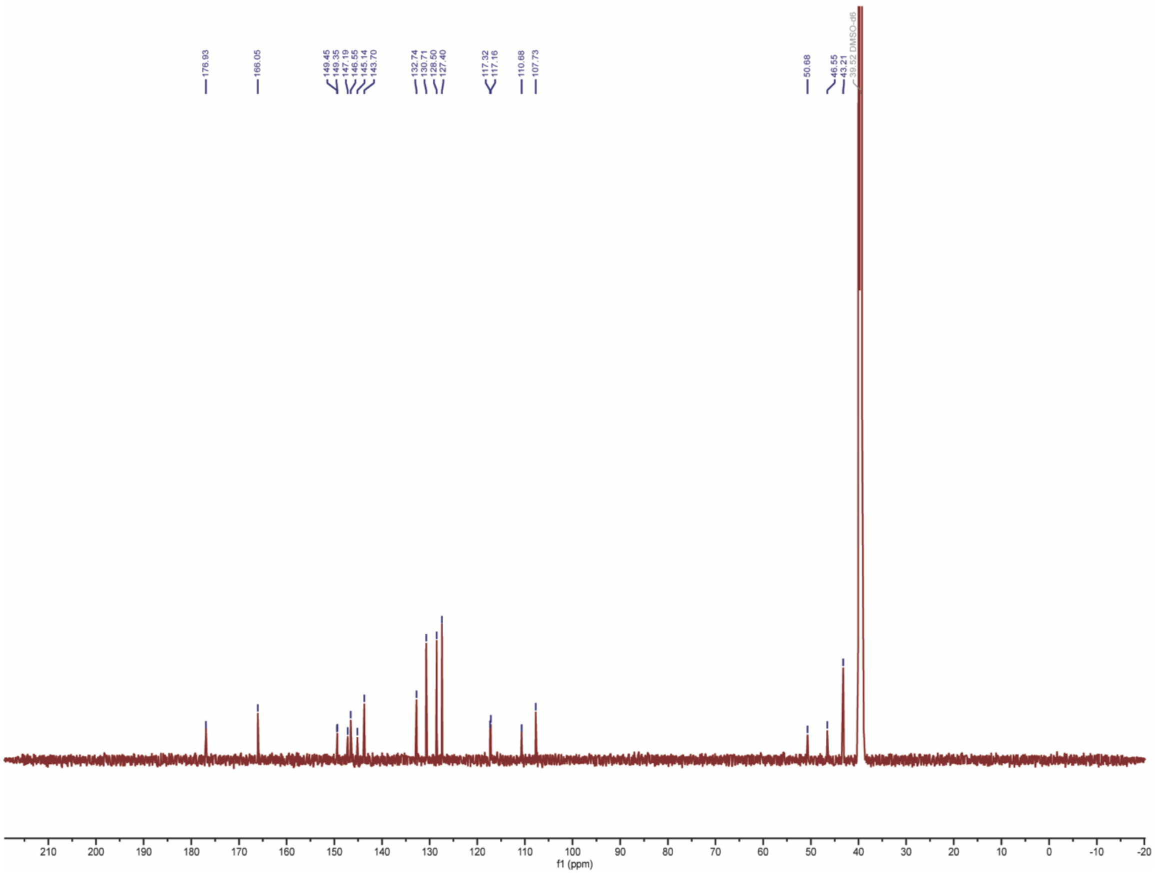
176.93
166.05
149.45
149.35
147.19
146.55
145.14
143.70
132.74
130.71
128.50
127.40
117.32
117.16
110.68
107.73
50.68
46.55
43.21
39.52 DMSO-d6
f1 (ppm)

### Bacterial Conditions to Generate a Resistance Strain

*Bacterial Culture Conditions*. *S. aureus* ATCC® 12600™ was cultured in cation-adjusted Mueller-Hinton broth at 37°C with shaking at 150 rpm. The antibiotic-resistant strain was cultured under the same conditions as its wild-type (WT) parent strain.

*Antimicrobial Susceptibility Testing*. Minimum inhibitory concentrations (MICs) were determined using the broth microdilution method, following Clinical and Laboratory Standards Institute (CLSI) standards and guidelines, with minimal modifications. Serial two-fold dilutions of compounds were prepared in sterile, clear round-bottom 96-well plates. To prepare microdilution trays, two-fold dilutions of antimicrobial agent were prepared in growth medium: Cation-adjusted Mueller-Hinton Broth (CAMHB) by adding 200 µL of the highest concentration to be tested (128 µg/mL, for example) in row A, mixing and transferring 100 µL from row A to 100 µL growth medium in row B, then repeating the mixing and transferring through row H of the 96-well plate, discarding the excess 100 µL remaining. This slight modification to the CLSI protocol enables evaluation of MICs for 3 compounds per plate in triplicate, albeit with only 8 compound dilutions (CLSI protocol enables 2 compounds in triplicate with 10 dilutions). Bacterial suspensions are added to a final concentration of $5 \times 10^5$ CFU/well by adding 5 µL of a 1:10 dilution of a 0.5 McFarland suspension ($1 \times 10^8$ CFU/mL) for each bacterium evaluated. Bacterial suspensions were prepared using the growth method described by CLSI. Well-isolated colonies (1 from an agar plate) were selected using a sterile loop and used to inoculate a tube containing 4 mL of CAMHB. The cultures are incubated at 37 ± 2° C until they achieve or exceed a turbidity of the 0.5 McFarland standard, determined by measuring OD600 (usually two to six hours). When growth exceeds a 0.5 McFarland standard, the turbidity is adjusted with broth to be equivalent to a 0.5 McFarland standard. Plates were incubated at 37 °C for 18-24 h, and the MIC was defined as the lowest concentration with no visible bacterial growth. OD600 readings were taken using a BioTek Synergy HTX multi-mode plate reader to quantify bacterial growth.

*Development of Resistant Strains*. Resistant strains were generated by culturing each bacterial strain on agar plates containing increasing concentrations of the antibacterial compounds, ranging from 100x to 400x the MIC values. Bacterial inoculum, ranging from $2.0 \times 10^9$ CFU/mL to $2.0 \times 10^1$ CFU/mL, was applied to each antibiotic-containing agar plate. This allowed a variety of antibiotic concentrations as well as bacterial concentrations to be studied, to facilitate the growth of resistant bacteria. Resistance colonies were isolated and purified by repeated plating. Confirmation of resistance was achieved by performing MIC assays and comparing the phenotypic values of the resistant strains to those of their wild-type counterparts (see Antimicrobial Susceptibility Testing for assay details).

### Sequencing Experiments

**RNA Isolation and Sequencing**. Total RNA was extracted from the bacterial strains using the Zymogen RNA extraction kit following the manufacturer's protocol. The RNA purity was determined by the A260/A280 ratio being ~2, and the integrity was visualized on a 0.5% agarose gel (Figure S2). The purified RNA was subject to 3`-poly-A tailing using a poly-A polymerase tailing kit from LGC Biosearch Technologies according to the manufacturer's protocol. The poly-A-tailed RNA was purified using a PCR cleanup kit and then quantified with a Qubit spectrometer. The end-prepared RNA was the input for the RNA direct nanopore sequencing library preparation kit from ONT (SQK-RNA004), following the manufacturer's protocol with one change to the method. The Induro (NEB) reverse transcriptase was used for the reverse transcription step with an extension time of 15 min at 55 °C, followed by denaturing the polymerase at 70 °C for 20 min. The library-prepared RNA was sequenced using the RNA flow cell (ONT) with the default parameters to generate passed reads with Q > 8.

The passed ONT reads were base-called with Dorado v0.5.2 via fast mode or hac-mode using the modification-aware models (v5.1.0) for Ψ, $m^5C$, or $m^6A$ (https://github.com/nanoporetech/dorado). The sequencing reads were aligned to the reference sequences using the align function in Dorado with default parameters. The reads were filtered by length using Samtools [1], followed by a second filtering step based on the 23S U1732C sequence mutation using pysam functions in Python v3.12. The modification-aware base calls were analyzed with modkit (https://github.com/nanoporetech/modkit) and visualized in IGV.[2] The extent of modification at each site was quantified from data with read depth >300 by adding to the high-probability (p>0.95) modification-aware base call frequencies the miscalls and indels for each site, as previously reported.[3] A student's t-test was conducted on replicate samples to determine statistical significance in RNA modification levels between the samples.

**DNA isolation and sequencing**. The genomic DNA was extracted from the bacterial strains using the Zymogen DNA extraction kit following the manufacturer's protocol. The DNA purity was determined by the A260/A280 ratio being ~1.8. Native DNA sequencing was performed with the extracted genomic DNA using the ligation sequencing kit v14 (ONT SQK-LSK114) following the manufacturer's protocol. As a note, the genomic DNA was not sheared to obtain long sequencing reads. The library-prepared genomic DNA was sequenced on a minION R10.4.1 flow cell (ONT) using default parameters to generate passed reads with Q > 10. The quality-passed reads were base called with Dorado (v0.5.2) using the sup model (v5.0.0). The base called data were aligned to the reference using Dorado with default settings. The aligned reads in BAM file format were sorted and indexed with Samtools and visualized in IGV.

**Figure S1**. Short-read sequencing mutants found in the *Sa*_A-692345$^{R}$ strain using the GATK pipeline.

| CHROM | POS | ID | REF | ALT | QUAL | DP |
|---|---|---|---|---|---|---|
| e1531a8e120f42ee_1 | | | 458101 | . | G | T |
| e1531a8e120f42ee_1 | | | 460115 | . | C | T |
| e1531a8e120f42ee_1 | | | 506240 | . | T | C |
| e1531a8e120f42ee_1 | | | 857675 | . | A | C |
| e1531a8e120f42ee_1 | | | 1039461 | . | AAAG | A |
| e1531a8e120f42ee_1 | | | 2027369 | . | AT | A |
| e1531a8e120f42ee_1 | | | 2059745 | . | T | C |
| e1531a8e120f42ee_1 | | | 2195100 | . | T | C |
| e1531a8e120f42ee_1 | | | 2505252 | . | C | T |
| e1531a8e120f42ee_1 | | | 2505337 | . | A | G |
| e1531a8e120f42ee_1 | | | 2505613 | . | A | C |
| e1531a8e120f42ee_1 | | | 2505636 | . | C | T |

The variant at position 506240 is 23S T1732C.

**Figure S2**. Agarose gel electrophoresis analysis of RNA directly sequenced.

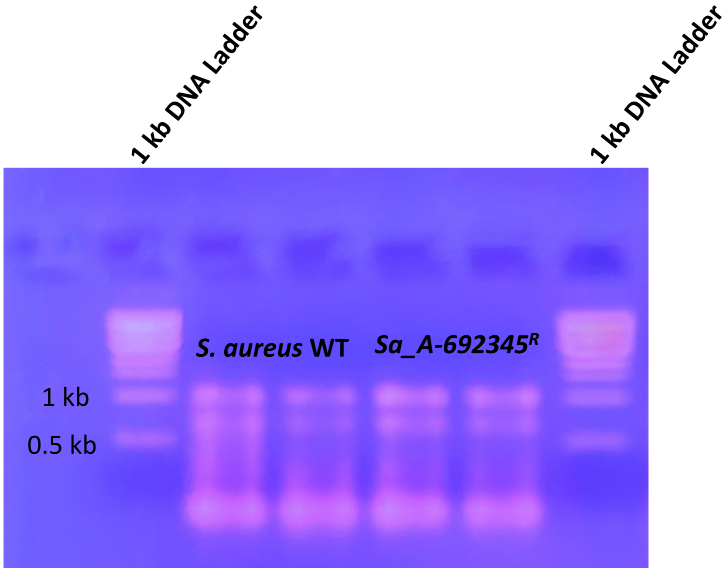


Agarose gel electrophoresis analysis was conducted to verify the integrity of the total RNA extracted from the *S. aureus* cells. The commercial ladder used for comparison was a 1 kb duplex DNA ladder, not an ssRNA ladder. In our hands, by the time the RNA ladders were received in the lab, they had already degraded to a point that rendered them unusable for comparison. The high stability of DNA is superior for these ladders; moreover, the gel was used to determine if the band profile was the same between the samples, which it was, and was never used for the estimation of the strand length.

**Figure S3**. RNA direct nanopore sequencing data visualized by IGV plots.

The RNA direct nanopore data were visualized in IGV. The visualizations were constructed using the default parameters in IGV. The color code for the plots is grey = >80% consensus between the reads and the reference nucleotide, and when the consensus is not met, A = green, C = blue, G = gold, T/U = red, and white space = indels.

16S rRNA: Top panel = WT *S. aureus*, bottom panel = *Sa*_A692345$^{R}$ strain

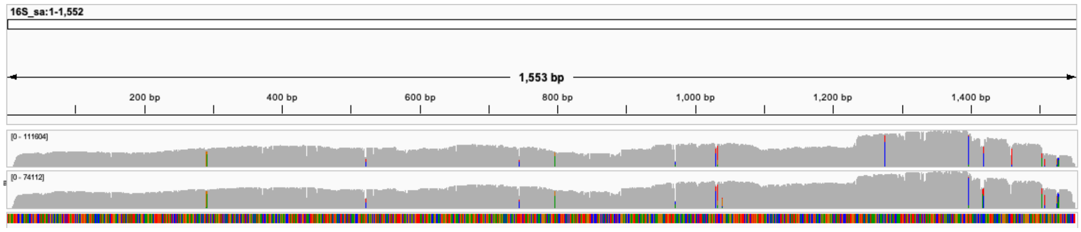


23S rRNA: Top panel = WT *S. aureus*, bottom panel = *Sa*_A692345$^{R}$ strain

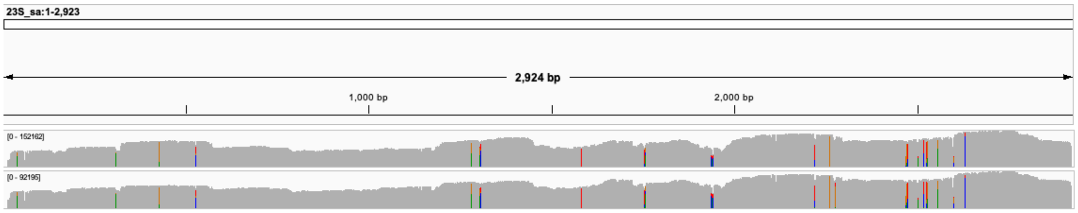


Filtered 23S rRNA U1732C sequence variants for the *Sa*_A692345$^{R}$ strain.

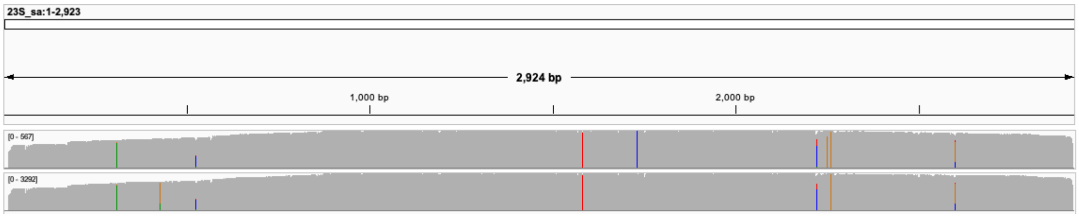


The C mutant is in the top panel, and U native sequence is in the bottom panel.

**Figure S4**. Comparisons of *S. aureus* and *E. coli* RNA direct nanopore data.

#### Tables Summarizing the rRNA Modification Findings

Green boxes represent the modification that was found, and white boxes represent the modification was not observed. The nanopore data are compared against two *S. aureus* cryo-EM structures solved in different strains 12600™ (Gonzalez-Lopez 2000)[4] or RN6390 (Golubev).[5]

| 16S rRNA Mod | *E. coli* Position | *S. aureus* Position | Cryo-EM (Strain NCTC 8325) Gonzalez-Lopez 2024 | Cryo-EM (Strain RN 6390) Golubev 2020 | Nanopore (Strain 12600) Present Report |
|---|---|---|---|---|---|
| Psi | 516 | 523 | | | |
| m7G | 527 | 537 | | | |
| m2G | 966 | 974 | | | |
| m5C | 967 | 975 | | | |
| m2G | 1207 | C1216 | | | |
| m4Cm | 1402 | 1411 | | | |
| m5C | 1407 | 1416 | | | |
| m3U | 1498 | 1508 | | | |
| m2G | 1516 | 1526 | | | |
| m6,6A | 1518 | 1528 | | | |
| M6,6A | 1519 | 1529 | | | |

| 23S rRNA Mod | *E. coli* Position | *S. aureus* Position | Cryo-EM (Strain NCTC 8325) Gonzalez-Lopez 2024 | Cryo-EM (Strain RN 6390) Golubev 2020 | Nanopore (Strain 12600) Present Report |
|---|---|---|---|---|---|
| m1G | 745 | 790 | | | |
| Psi | 746 | 791 | | | |
| m5U | 747 | 792 | | | |
| Psi | 955 | 999 | | | |
| m6A | 1618 | 1662 | | | |
| m2G | 1835 | 1862 | | | |
| Psi | 1911 | 1938 | | | |
| m3Psi | 1915 | 1942 | | | |
| Psi | 1917 | 1944 | | | |
| Cm | 1920 | 1947 | | | |
| m5U | 1939 | 1966 | | | |
| m5C | 1962 | 1989 | | | |
| m6A | 2030 | 2057 | | | |
| m7G | 2069 | 2096 | | | |
| Gm | 2251 | 2278 | | | |
| m2G | 2445 | 2472 | | | |

| 23S rRNA Mod | *E. coli* Position | *S. aureus* Position | Cryo-EM (Strain NCTC 8325) Gonzalez-Lopez 2024 | Cryo-EM (Strain RN 6390) Golubev 2020 | Nanopore (Strain 12600) Present Report |
|---|---|---|---|---|---|
| D | 2449 | 2476 | | | |
| Psi | 2457 | 2484 | | | |
| Cm | 2498 | 2525 | | | |
| ho5C | 2501 | 2528 | | | |
| m2A | 2503 | 2530 | | | |
| Psi | 2504 | 2531 | | | |
| Um | 2552 | 2579 | | | |
| Psi | 2580 | 2607 | | | |
| Psi | 2604 | 2631 | | | |
| Psi | 2605 | 2632 | | | |

The RNA direct nanopore data visualized in IGV are laid out in the following order. The top block of sequencing tracks provides data obtained from *E. coli*. The rRNA modifications are marked using *E. coli* numbering. The bottom block of sequencing tracks provides data for *S. aureus*. In each track in the blocks from top to bottom, the first track is data obtained without using a modification-aware model, in which grey => 80% consensus between the reads and the reference nucleotide, and when the consensus is not met, A = green, C = blue, G = gold, T/U = red, and white space = indels. The second track provides data using the m$^6$A model, with m$^6$A calls shown in purple. The third track provides data using the m$^5$C model, with m$^5$C calls shown in pink. The fourth track provides data using the Ψ model, with Ψ calls shown in aqua. Below each figure is commentary regarding data interpretation and comparison to the two reported cryo-EM structures for the *S. aureus* ribosome.

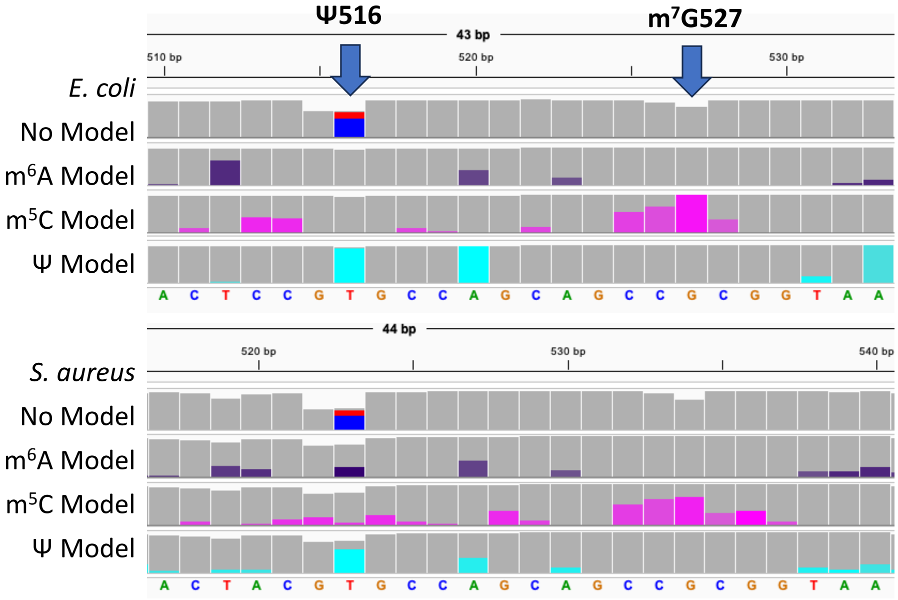


The *E. coli* 16S rRNA Ψ516 is found in *S. aureus* at position 523 based on the base miscall in the track without a model and in the Ψ-aware base call model for each organism. This site was not observed by cryo-EM analyses of the *S. aureus* ribosome (Golubev 2020 and Gonzalez-Lopez 2024). The *E. coli* m$^7$G527 was observed in *S. aures* in both cryo-EM structures (Golubev 2020 and Gonzalez-Lopez 2024), and the nanopore data identify this modification based on the miscall analysis via increased indels at the site.

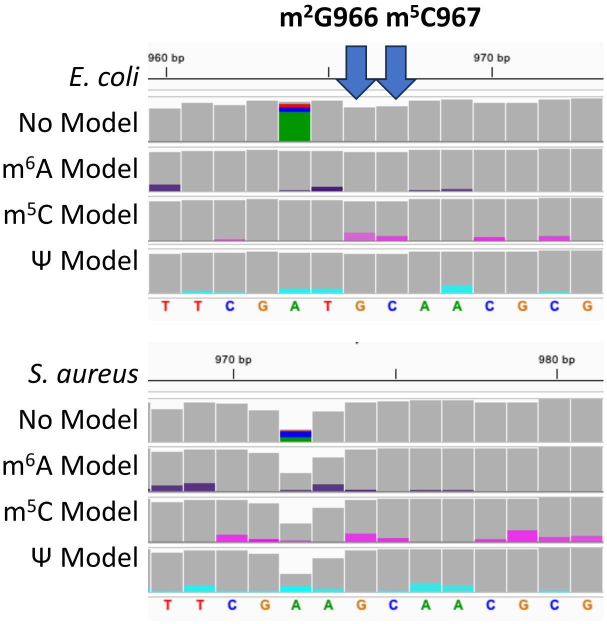


The *E. coli* 16S rRNA m$^2$G966 and m$^5$C967 modifications were observed in the *S. aureus* in both cryo-EM structures of the ribosome (Golubev 2020 and Gonzalez-Lopez 2024). In the nanopore data for *S. aureus* 16S rRNA, the m$^2$G is observed by an increase in base call error, and the m$^5$C is observed by a comparable signature in the m$^5$C-aware base call model as observed in the *E. coli* rRNA.

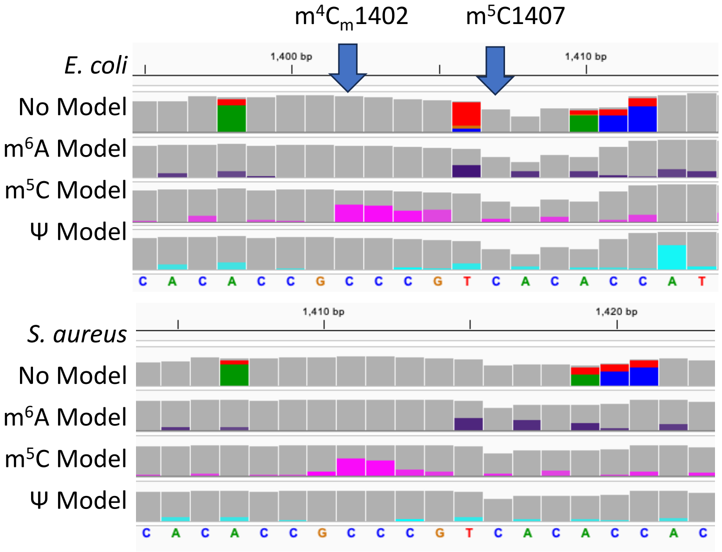


The *E. coli* 16S rRNA m$^4$C$_m$1402 and m$^5$C1407 modifications were observed in the *S. aureus* at positions 1411 and 1413, respectively, in the nanopore data based on the m$^5$C-aware base call data and miscall analysis. The m$^4$C$_m$1402 was found in both cryo-EM structures of the *S. aureus* ribosome (Golubev 2020 and Gonzalez-Lopez 2024).

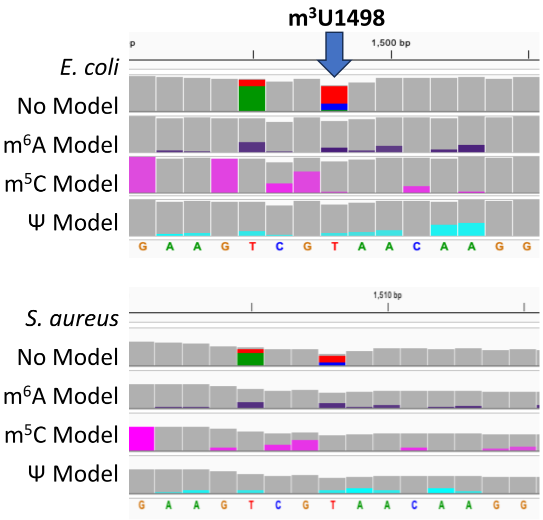


The *E. coli* 16S rRNA $m^5U1498$ was observed in the *S. aureus* rRNA based on base miscall analysis. This modification was also observed in both cryo-EM structures of the *S. aureus* ribosome (Golubev 2020 and Gonzalez-Lopez 2024).

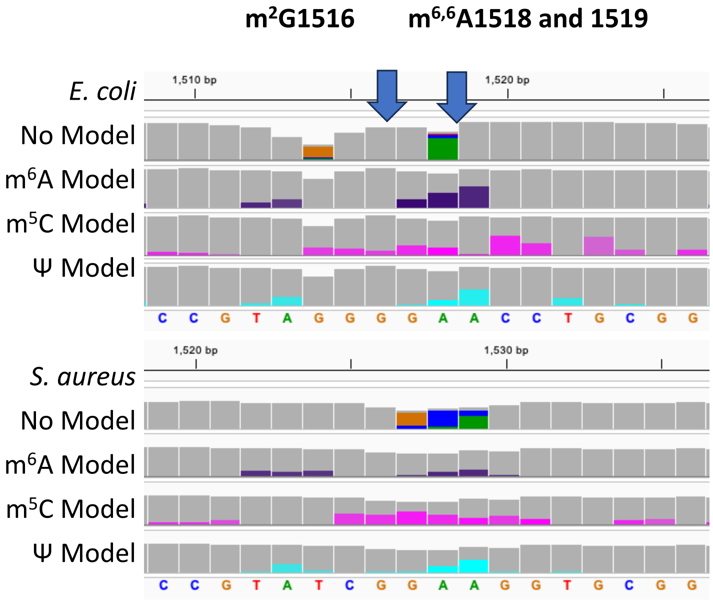


The *E. coli* 16S rRNA $m^2G1516$ was not observed in the S. aureus rRNA, and the two $m^{6,6}A$ residues at 1518 and 1519 were observed in the S. aureus rRNA based on base miscall analysis and $m^6A$-aware base calling. The two $m^{6,6}A$ residues were observed in both cryo-EM structures of the *S. aureus* ribosome (Golubev 2020 and Gonzalez-Lopez 2024).

**m[1]G745, Ψ746, and m[5]U747**

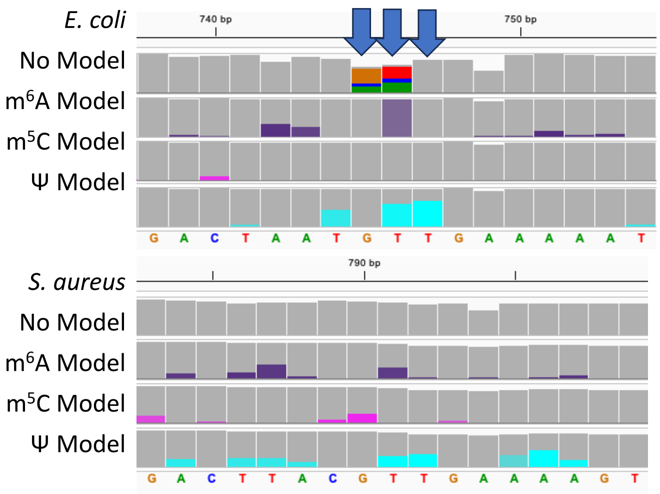


The *E. coli* 23S rRNA m[1]G745 was not observed in the *S. aureus* rRNA, while the Ψ746 and m[5]U747 were observed in the S. aureus rRNA based on base miscall analysis and Ψ-aware base calling. The Ψ791 in *S. aureus* is now identified by nanopore sequencing. The m[5]U792 in *S. aureus* was observed in one cryo-EM structure of the ribosome (Gonzalez-Lopez 2024).

**m[6]A1618**

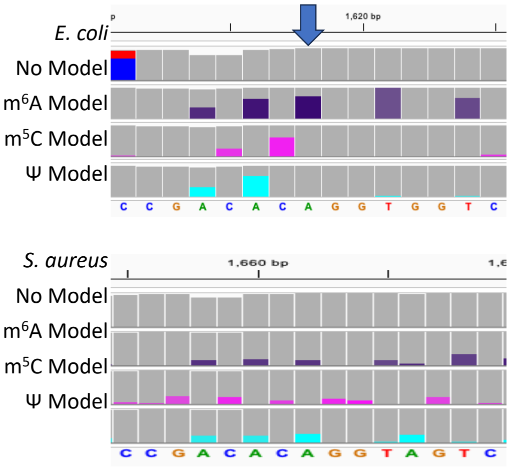


**m[6]A2030**

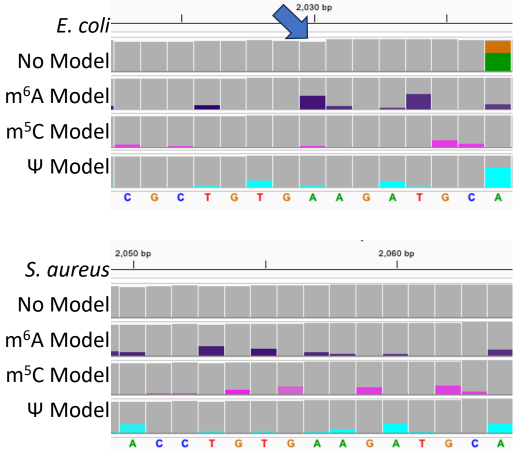


The *E. coli* 23S rRNA has two m[6]A residues at positions 1618 and 2030. Neither of the corresponding residues in S. aureus appears to be modified based on m[6]A-aware base calling. Neither of these sites was reported to be modified in the cryo-EM structures (Golubev 2020 and Gonzalez-Lopez 2024).

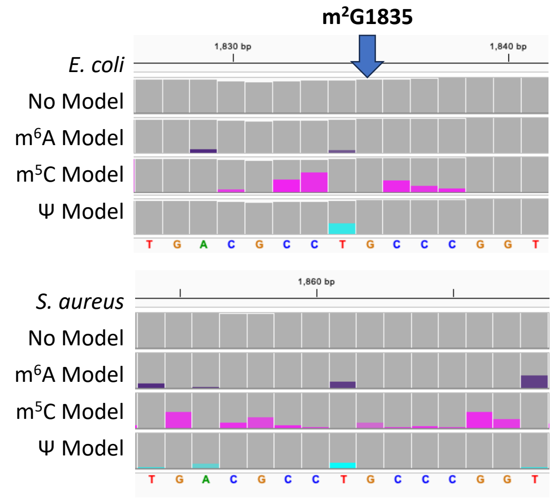


The *E. coli* 23S rRNA m$^{2}$G at position 1835 was not observed in the nanopore data or the two cryo-EM structures (Golubev 2020 and Gonzalez-Lopez 2024).

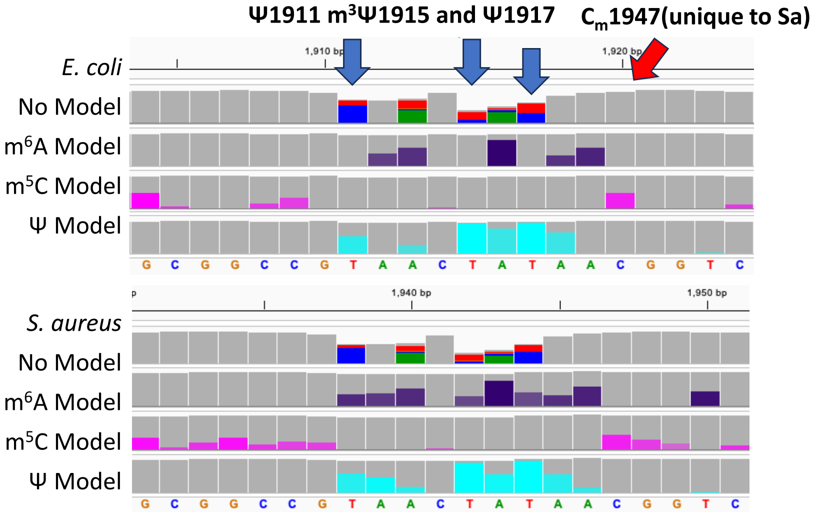


In the 23S rRNA of *E. coli* are the modifications Ψ1911, m$^{3}$Ψ1915, and Ψ1917. All three Ψ modifications were observed in *S. aureus* at positions 1938, 1942, and 1944 in the nanopore data. The methylation status of 1942 cannot be determined by the nanopore data; however, this was the only modification observed in one of the cryo-EM structures of the *S. aureus* ribosome (Gonzalez-Lopez 2024), supporting it is m$^{3}$Ψ in the present analysis. The *S. aureus* ribosome possesses Cm1947 (S. aureus numbering) that is not observed in *E. coli*. The C$_{m}$1947 was observed in one cryo-EM structure (Golubev 2020), and the nanopore data support its presence on the basis of m$^{5}$C-aware base call data analysis.

**$m^5U1939$**

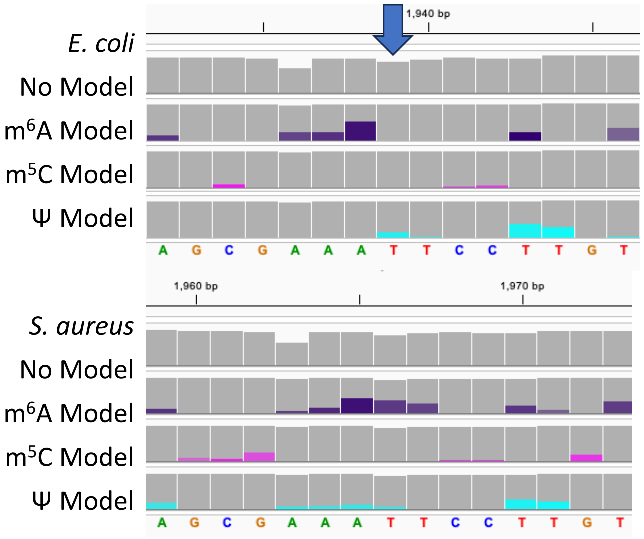


In the 23S rRNA of *E. coli* is $m^5U1939$, which was observed by one cryo-EM structure of the S. aureus ribosome at position 1966 (Golubev 2020); however, the nanopore data do not provide a signature for this modification.

**$m^5C1962$**

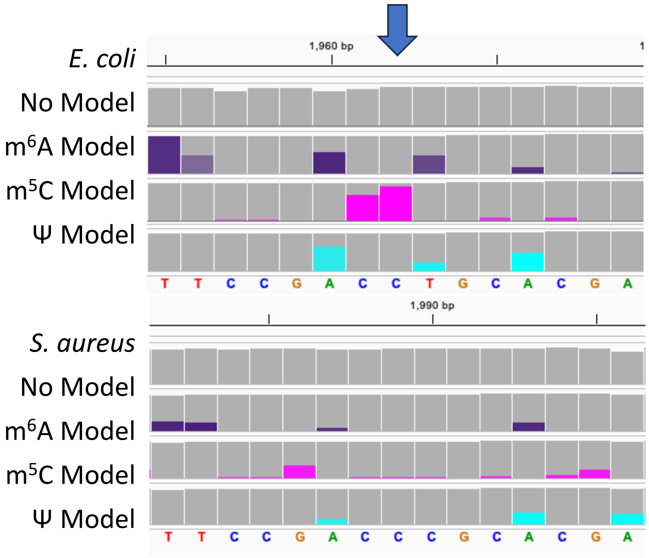


In the 23S rRNA of *E. coli* is $m^5C1962$, which failed to be observed by nanopore sequencing and in both cryo-EM structures.

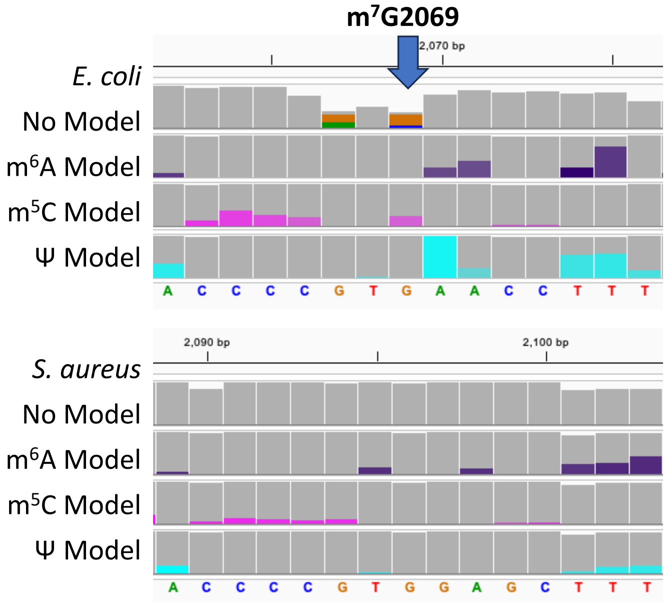


In the 23S rRNA of *E. coli* is m$^7$G2069, which was not observed by nanopore sequencing and in both cryo-EM structures.

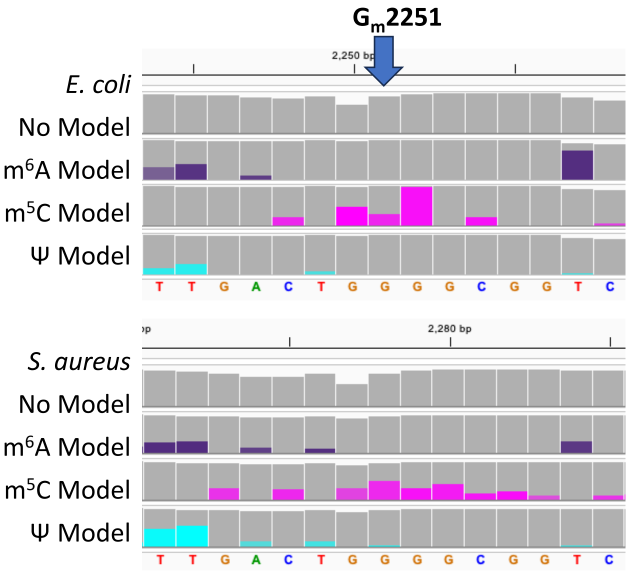


In the 23S rRNA of *E. coli* is G$_m$2251, it was observed at position 2278 in one cryo-EM structure of the *S. aureus* ribosome and in the nanopore sequencing data as an increased base miscall signature.

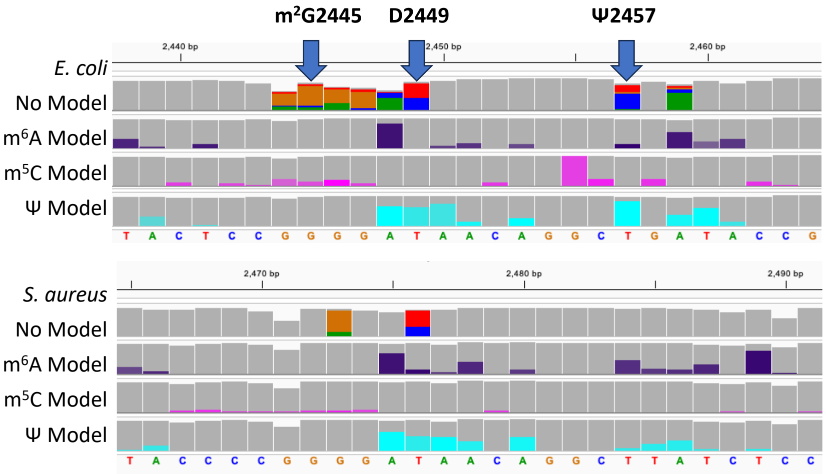


In the 23S rRNA of *E. coli* are the modifications m$^2$G2445, D2449, and Ψ2457. The nanopore sequencing data support the presence of D at position 2476 in the *S. aureus* 23S rRNA, which is supported by its presence in one cryo-EM structure (Gonzalez-Lopez 2024). In *S. aureus,* m$^2$G2472 was observed in a cryo-EM structure of the ribosome (Golubev 2020), but it was not observed in the nanopore data. Lastly, there is no evidence that U2484 is a Ψ in *S. aureus* based on the nanopore Ψ-aware base call data.

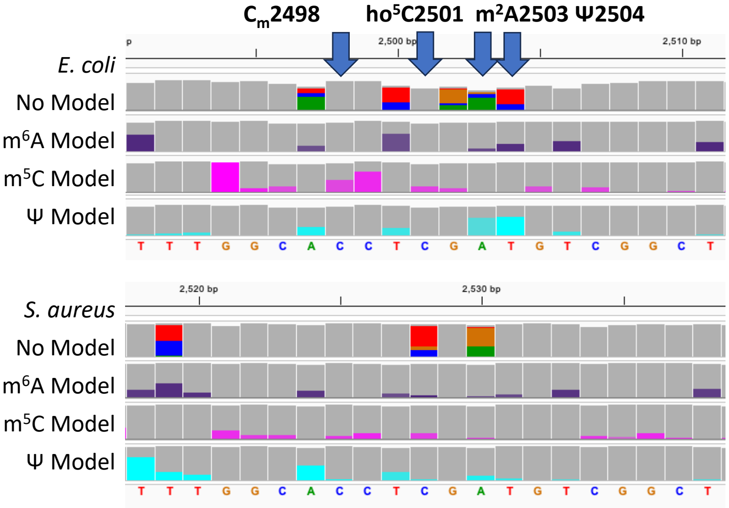


In the 23S rRNA of *E. coli* are the modifications $C_m$2498, ho$^5$C2501, m$^2$A2503, and Ψ2504. The nanopore data support the presence of ho$^5$C and m$^2$A at positions 2528 and 2530 in the *S. aureus* 23S rRNA, respectively. The m$^2$A2530 modification was observed in both cryo-EM structures for the *S. aureus* ribosome (Golubev 2020 and Gonzalez-Lopez 2024). The nanopore data suggest U2531 is not a Ψ in *S. aureus*.

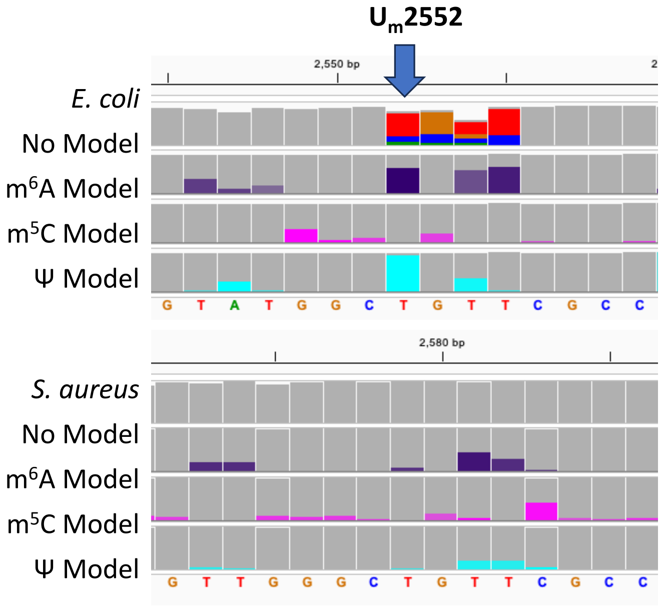


In the 23S rRNA of *E. coli* is $U_m$2552. There is no evidence that the corresponding U in *S. aureus*, 2579, is modified based on the nanopore data. Additionally, this site was not observed to be modified in either cryo-EM structure of the *S. aureus* ribosome.

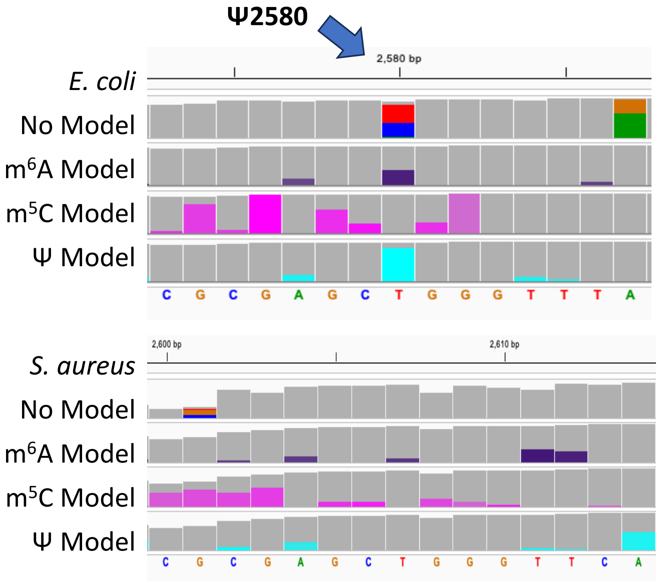


In the 23S rRNA of *E. coli* resides a Ψ at position 2580. The nanopore and cryo-EM data support the corresponding position in S. aureus (U2607) is not modified.

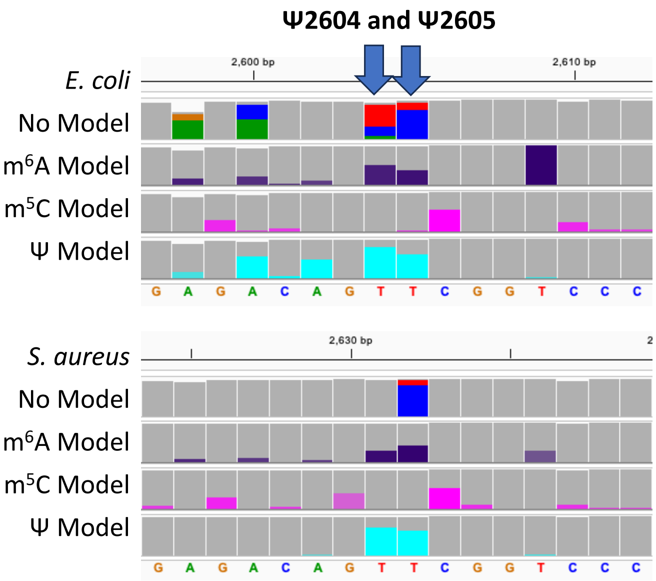


In the 23S rRNA of *E. coli* resides two Ψ residues at positions 2604 and 2605. The nanopore data via Ψ-aware base calling support in the 23S rRNA of *S. aureus,* there are two Ψ residues at positions 2631 and 2632.
